# Classifying cell cycle states and a quiescent-like G0 state using single-cell transcriptomics

**DOI:** 10.1101/2024.04.16.589816

**Authors:** Samantha A. O‘Connor, Leonor Garcia, Rori Hoover, Anoop P. Patel, Benjamin B. Bartelle, Jean-Philippe Hugnot, Patrick J. Paddison, Christopher L. Plaisier

## Abstract

Single-cell transcriptomics has unveiled a vast landscape of cellular heterogeneity in which the cell cycle is a significant component. We trained a high-resolution cell cycle classifier (ccAFv2) using single cell RNA-seq (scRNA-seq) characterized human neural stem cells. The ccAFv2 classifies six cell cycle states (G1, Late G1, S, S/G2, G2/M, and M/Early G1) and a quiescent-like G0 state (Neural G0), and it incorporates a tunable parameter to filter out less certain classifications. The ccAFv2 classifier performed better than or equivalent to other state-of-the-art methods even while classifying more cell cycle states, including G0. We demonstrate that the ccAFv2 classifier effectively generalizes the S, S/G2, G2/M, and M/Early G1 states across cell types derived from all three germ layers. While the G0, G1, and Late G1 states perform well in neuroepithelial cell types, their accuracy is lower in other cell types. However, misclassifications are confined to the G0, G1, and Late G1 states. We showcased the versatility of ccAFv2 by successfully applying it to classify cells, nuclei, and spatial transcriptomics data in humans and mice, using various normalization methods and gene identifiers. We provide methods to regress the cell cycle expression patterns out of single cell or nuclei data to uncover underlying biological signals. The classifier can be used either as an R package integrated with Seurat or a PyPI package integrated with SCANPY. We proved that ccAFv2 has enhanced accuracy, flexibility, and adaptability across various experimental conditions, establishing ccAFv2 as a powerful tool for dissecting complex biological systems, unraveling cellular heterogeneity, and deciphering the molecular mechanisms by which proliferation and quiescence affect cellular processes.

## Introduction

Single-cell RNA sequencing (scRNA-seq) is a robust method for dissecting the transcriptional states of individual cells obtained from specific conditions. These cellular transcriptional states are influenced by various biological signals, including cell type and the phase of the cell cycle. The cell cycle is a tightly regulated and intricately coordinated biological process that orchestrates the division of a cell into two daughter cells. Adult stem cell populations often reside in a quiescent G0 state outside of the cell cycle, reactivating only upon receiving appropriate signals to divide (Doetsch 2003; Obernier et al. 2018). Current state of the art methods to predict cell cycle states based on scRNA-seq transcriptome profiles lump G0 cells with G1 cells (Hao et al. 2021; Zheng et al. 2022; Schwabe et al. 2020; Liu et al. 2017; Hsiao et al. 2020; Scialdone et al. 2015). The grouping of G0 with G1 fails to recognize the clear differences in expression patterns and quiescent phenotype displayed by G0 cells, making them readily distinguishable from G1 cells (O’Connor et al. 2021). The aim of this research is to develop a cell cycle classifier capable of identifying the G0 state in neuroepithelial cells and to determine whether this state can be generalized to other cell types.

Developing cell cycle classifiers from scRNA-seq transcriptional profiles is challenging due to the scarcity of datasets with experimentally validated ground truth cell cycle labels. Training a classifier requires having example transcriptome profiles labeled with cell cycle states. Previous studies have used Hoechst (DNA stain; (Buettner et al. 2015) or FUCCI (Leng et al. 2015) to sort embryonic stem cells into G1, S, and G2M subpopulations. However, there are some caveats to these studies. Firstly, these cells were not fixed, meaning they could continue to cycle after sorting and may not be transcriptionally in the same state as they were when they were sorted. Secondly, the markers used for sorting focused on DNA, protein, and post-translational modification abundances, which may not accurately reflect the transcriptional state of the cells. Thirdly, it has been established that embryonic stem cells do not have well-defined G1 or G0 cell cycle states as they quickly transition through cell cycles to produce many cells in the embryo (Ballabeni et al. 2011; White and Dalton 2005). In preliminary analyses it was found that the cell cycle labels were significantly out of alignment (error rates ≥ 0.7) with the transcription states of the cells as determined by ccSeurat (**Supplemental Table S1**), which is the *de facto* standard in the field.

Previously, we used scRNA-seq of U5 human Neural Stem Cells (U5-hNSCs; Davis and Temple 1994; Johe et al. 1996) grown *in vitro* to discern seven cell cycle states including a quiescent-like G0 state (O’Connor et al. 2021). An Artificial Neural Network (ANN) (Ma and Pellegrini 2020) classifier named the cell cycle ASU/Fred Hutch (ccAF) was trained to predict these seven cell cycle states in cells from new datasets (O’Connor et al. 2021). In those studies, the ccAF classifier was applied to a host of neuroepithelial derived cells characterized by scRNA-seq, including glioblastoma patient tumor cells. The underlying software packages for constructing ANNs (TensorFlow and Keras) have been significantly improved and we hypothesized that reimplementation of the ccAF classifier would significantly improve classifier performance and provide likelihoods for each classification, a feature not available in the original ccAF implementation.

In addition to the advancements in ANN methodology, numerous new scRNA-seq studies have been conducted that include actively dividing cells. Particularly valuable for assessing the quality and generalizability of the classifier is an atlas of 245,906 cells from 15 different cell types, spanning all three germ layers, derived from human fetal tissue 3 to 12 weeks post-conception (Zeng et al. 2023). A second atlas of developing human spinal cord (Zhang et al. 2021) will be used to evaluate whether the classifier can be applied to both single cell and single nuclei RNA-seq (scRNA-seq and snRNA-seq). An atlas of adult neurogenesis in the ventricular-subventricular zone (V-SVZ) (Cebrian-Silla et al. 2021) will be used to demonstrate that the classifier can be applied to mouse cells. It will also allow comparisons to be made between the cell cycle proportions of cell types from adult mouse neurogenesis in the V-SVZ (Cebrian-Silla et al. 2021) and the developing human telencephalon (Nowakowski et al. 2017). Two studies of quiescent neural stem cells will be crucial for demonstrating the identity of the G0 cell state (Llorens-Bobadilla et al. 2015; Dulken et al. 2017). Additionally, we collected scRNA-seq for two IDH mutant low-grade glioma (LGG) cell lines in conditions with and without growth factors. This will allow us to gain insights into the performance of the classifier when confronted with a higher proportion of non-cycling cells. We will also apply the classifier to *in vivo* glioblastoma tumor cells and *in vitro* glioblastoma tumor derived cancer stem cells that were not included in the previous ccAF classifier studies (Couturier et al. 2020). Finally, application of the classifier to a high-resolution spatial-transcriptomics (ST-seq) study of a mouse embryo at 15.5 weeks post-conception (E15.5) will allow us to resolve canonical biological and morphological phenomena for the developmental stage. These datasets offer a robust foundation for rigorously testing and validating the improved ccAF version 2 (ccAFv2) classifier, showcasing its versatility across species, single cells and nuclei, and generalizability across cell types from all germ layers.

The goal of this research is to develop an improved cell cycle classifier using current state of the art machine learning technology. We aim to demonstrate that the classifier outperforms existing models and generalizes well across various cell types, library preparation methods (scRNA-seq, snRNA-seq, ST-seq), gene annotations, and normalization techniques. Lastly, we aim to provide a classifier with a more user-friendly interface to facilitate its application in future studies.

## Results

### Implementation of neural network classifier for ccAFv2

We implemented the core algorithm of ccAFv2 to take advantage of significant improvements in machine learning tools that should improve classifier performance and provide likelihoods for each predicted cell cycle classification. The ccAFv2 core algorithm is broken up into two steps. <u>First</u>, the input data is run through the artificial neural network (ANN) to compute likelihoods for each class (i.e., Neural G0, G1, Late G1, S, S/G2, G2/M or M/Early G1; **Figure 1A-B**). The underlying ANN for ccAFv2 starts with a dense input layer connected to two hidden layers that connect to a softmax output layer (**Figure 1A**). Overfitting in the ANN is mitigated by dropout regularization via two dropout layers. The first dropout layer is positioned between the first and second hidden layers and the second dropout layer is between the second hidden layer and the softmax output (**Figure 1A**; Xie et al. 2019). <u>Second</u>, the likelihoods calculated by the ANN for each cell cycle state are used to determine which state should be assigned for each cell (**Figure 1B**). The cell cycle state with the maximum likelihood is identified and if the likelihood is greater than or equal to the likelihood threshold then the state is returned. Otherwise, if the maximum likelihood is less than the likelihood threshold a state of “Unknown” is returned. These improvements to the core ANN of ccAFv2 will be rigorously tested in the subsequent sections.

**Figure 1.**
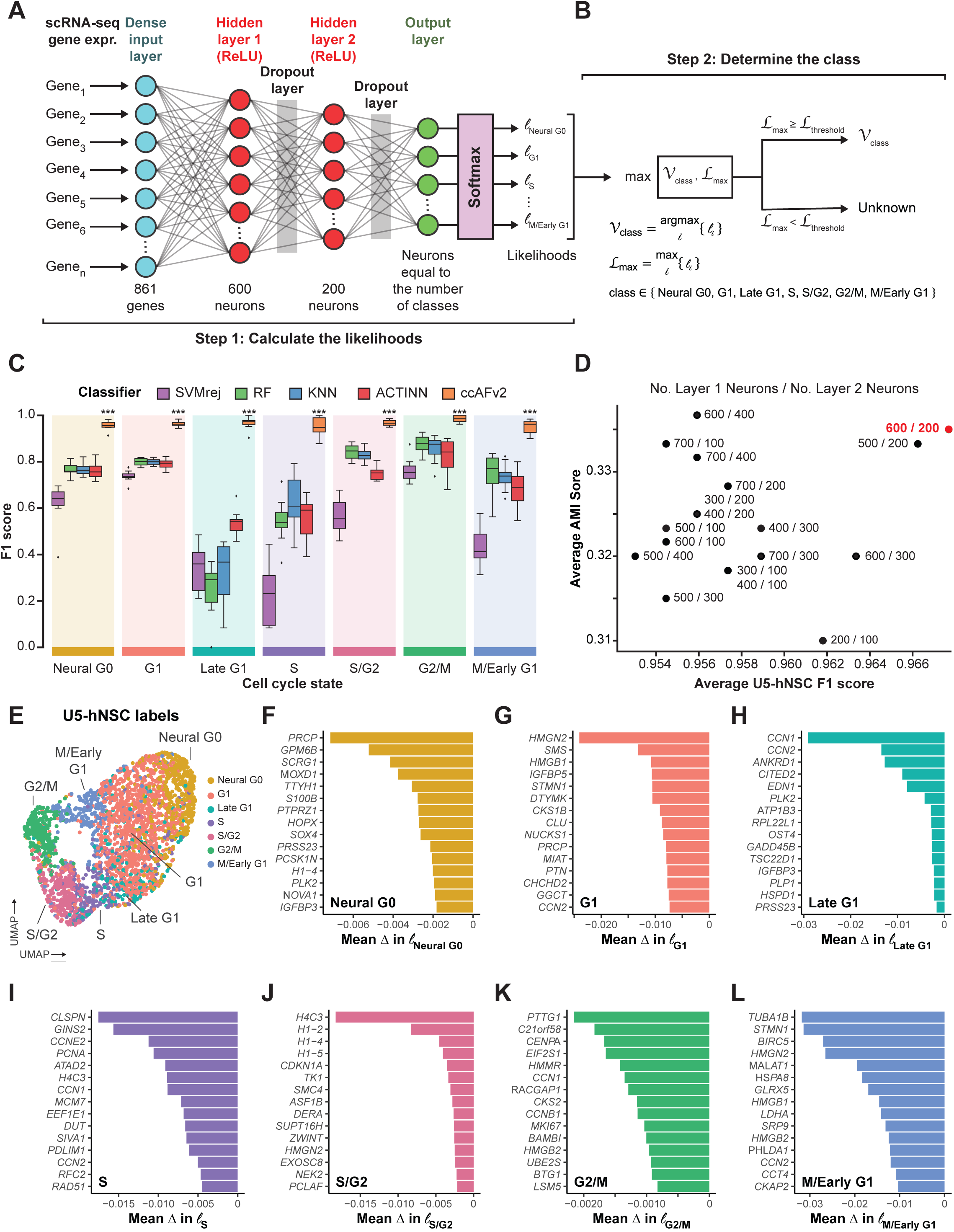
Implementing and testing the ccAFv2 classifier. **A.** The design of the Artificial Neural Network (ANN) implemented for the ccAFv2. Expr. = expression, ReLU = Rectified Linear Units. **B.** Method designed to determine the predicted class from the likelihoods generated by running expression data from a single cell through the ccAFv2 ANN. **C.** Comparison of five different classification methods using F1 scores (a metric that integrates precision and recall, and has a maximum value of 1), from the 10-fold cross validation analysis of training on the U5-hNSCs. The F1 scores are computed for each cell cycle state from each of the 10 testing datasets. **D.** Determining the optimal number of neurons in each hidden layer using average U5-hNSC F1 score across cell cycle states on the x-axis, and the average AMI score across the remaining datasets (U5-hNSCs; glioma stem cells: BT322, BT324, BT326, BT333, BT363, BT368; tumor cells: BT363, BT368; and Grade 2 Astrocytoma: LGG275). Each combination of hidden layer neurons is labeled using: number of hidden layer one neurons / number of hidden layer two neurons. The chosen optimal configuration of 600 hidden layer 1 neurons and 200 hidden layer 2 neurons (600 / 200) is denoted in red. **E.** UMAP of U5-hNSCs with cells colored by the labels from O’Connor et al., 2021. **F**-**L.** The top 15 most important features for the ccAFv2 classifier were identified based on the mean change (Δ) in likelihood after permuting each feature’s expression. A negative mean change in likelihood indicates that the feature increased the likelihood of predicting a ccAFv2 state.

### Training the ccAFv2 classifier

The training data for ccAFv2 is comprised of scRNA-seq from actively dividing U5 human neural stem cells (U5-hNSCs) cultured *in vitro* (O’Connor et al. 2021). The U5-hNSCs were cultivated from the telencephalon of a human fetus 8 weeks post-conception (Bressan et al. 2017). We previously identified 7 transcriptional states in the U5-hNSCs that were mapped to cell cycle states (i.e., Neural G0, G1, Late G1, S, S/G2, G2/M, and M/Early G1; O’Connor et al. 2021).

The U5-hNSC scRNA-seq data were reanalyzed using current quality control and normalization methods which resulted in 2,962 good quality single-cell transcriptome profiles (**Supplemental Figure S1A**). The U5-hNSC scRNA-seq profiles, along with the previously established cell cycle labels (O’Connor et al. 2021), represent the most meticulously curated training dataset available for cell cycle classification.

We compared the newly implemented ccAFv2 classifier against four distinct classification methods: support vector machine with rejection (SVMrej), random forest (RF), scRNA-seq optimized *k*-nearest neighbor (KNN), and ACTINN (Ma and Pellegrini 2020) which was used to build ccAF (O’Connor et al. 2021). The training dataset for all classifiers consisted of the pre-processed U5-hNSC scRNA-seq subset to the 861 genes upregulated in cell cycle states (log_2_FC ≥ 0.25, adjusted p-value ≤ 0.05; **Supplemental Table S2**). We applied 10-fold cross-validation (CV) for each classification method (**Supplemental Figure S1A**) and observed that ccAFv2 exhibited significantly improved F1 scores for each cell cycle state compared to other classification methods (p-values ≤ 2.8 x 10^-6^; **Figure 1C**), establishing it as the most accurate cell cycle classifier overall. A benefit of using the F1-score as the performance metric is that it accounts for the imbalance in class label proportions within the training set. We evaluated the impact of balancing label proportions in the training dataset, but this resulted in worse model performance (**Supplemental Figure S1B**). The accuracy of ccAFv2 when applied to U5-hNSCs was 88.4%, and the main difference when compared to ccAF was an improvement in Late G1 cell predictions (**Supplemental Figure S1C**-**D**). The overall error rate for ccAFv2 was 3.3%, which is a considerable improvement from the 18.4% of ccAF (O’Connor et al. 2021). The reimplementation of the ANN for the ccAFv2 classifier has significantly improved its performance across all cell cycle states, providing a robust foundation for further optimization and comprehensive characterization of its capabilities.

### Optimizing the number of neurons in hidden layers

A crucial factor in optimizing the parameters of the ccAFv2 ANN was determining the ideal number of neurons in each hidden layer. We conducted a systematic comparison of 18 different combinations for the number of neurons in the two hidden layers (first hidden layer: ranging from 200 to 700 neurons, and second hidden layer: ranging from 100 to 400 neurons) across U5-hNSCs (O’Connor et al. 2021), a low grade glioma stem cell line (LGG275), six glioma stem cell lines (BT322, BT324, BT326, BT333, BT363, and BT368; Couturier et al. 2020), and two glioma tumors (BT363 and BT368; Couturier et al. 2020). The optimal combination was determined by having the highest average F1-score and Adjusted Mutual Information (AMI) score using ccSeurat as the reference (**Figure 1D**; **Supplemental Table S3**). We chose to employ the ccSeurat classifier (Butler et al. 2018) to predict the reference labels because true cell cycle state labels do not exist for all datasets. The ccSeurat classifier was chosen for three reasons: 1) it is the de facto standard method for cell cycle classification currently, 2) it performs well when applied to many different datasets, and 3) it uses a totally different underlying algorithm to classify cell cycle state than ccAFv2. We found that configuring the ccAFv2 ANN with 600 neurons in the first hidden layer and 200 in the second hidden layer yielded the largest average F1 score and second largest AMI score (**Figure 1D**). This specific parameterization has been assigned for the hidden layers of the ccAFv2 ANN, and all prior and subsequent ccAFv2 classifications use this parameterization.

### Most important features for classifying ccAFv2 states

After optimizing the training of the ccAFv2 ANN, it is sensible to determine which features are most essential for classifying each of the seven states. We computed feature importance by permuting one of the 861 genes in the U5-hNSCs dataset and asking what impact that had on the likelihoods for each of the seven states. Randomizing the expression of an important feature for classifying a ccAFv2 state would lead to reductions in the states likelihood for cells known to be of this state. Thus, it is crucial that the dataset used for feature importance have cell cycle labels, which is why the U5-hNSCs were used for feature importance analyses (**Figure 1E**). We report the top 15 most important genes for each of seven ccAFv2 states (**Figure 1F-L**).

Eleven of the most important genes for classifying the Neural G0 state (**Figure 1F**) were also marker genes of Neural G0 in the U5-hNSCs. The first most important gene for classifying the G1 state (**Figure 1G**) was *HMGN2*, and in prior studies over-expression of *HMGN2* in osteosarcoma cells led to significantly higher number of cells in G0/G1 (Liang et al. 2015). The top two most important genes for the classifying the Late G1 state (**Figure 1H**) include two Immediate-Early Genes (IEGs) *CCN1* and *CCN2* which are known to be induced rapidly after initiation of cell cycle progression by many factors (Tullai et al. 2007). The top four most important genes for classifying the S state (**Figure 1I**) include three genes required for DNA replication during S phase (*CLSPN*, *GINS2*, and *PCNA*) and the cyclin associated with S phase (*CCNE2*). The top four most important genes for classifying the S/G2 state (**Figure 1J**) are all histones, specifically one *H4* histone and multiple *H1* histones isoforms that enable the condensation of nucleosomes into chromatin. The top five most important genes for classifying the G2/M state (**Figure 1K**) include a gene involved in keeping sister chromatids from separating (*PTTG1*), and two genes involved in kinetochore and centromere maintenance and function (*CENPA*, *HMMR*; Maxwell et al. 2005). Additionally, the ninth most important gene for classifying G2/M is *CCNB1* the cyclin that peaks in mitosis, and *MKI67* which is an established marker of cell proliferation (Scholzen and Gerdes 2000). Finally, the top three most important genes for classifying the M/Early G1 state (**Figure 1L**) are a microtubule component protein *TUBA1B*, a microtubule associated protein *STMN1*, and a component of the chromosome passage protein complex (CPC) which is essential for sister chromatid alignment and segregation during mitosis and cytokinesis (Vong et al. 2005). The functions of the key genes for classifying each state align well with the molecular processes of each cell cycle state, supporting the conclusion that the identified classes in U5-hNSCs reflect the underlying biology of the cell cycle.

Next, we evaluated the expression of important genes for each ccAFv2 state in an independent dataset, the *in vivo* hNSCs collected from whole fetal brain at 9 weeks post-conception (PCW 9 R1; Zeng et al. 2023). This allows us to assess the generalizability of these genes as key markers across novel datasets, providing insight into their broader applicability and robustness. The expression of important genes for all ccAFv2 states were expressed strongly in the state they marked, except for the important genes for the G1 state (**Supplemental Figure S1E**). In our prior study it was difficult to identify markers for G1 phase cells, and so the lack of translation for important genes for the G1 state is not surprising. The successful translation of key genes to an independent dataset supports the hypothesis that ccAFv2 and the marker genes identified in U5-hNSCs are broadly applicable to *in vivo* hNSCs.

### Comparison with existing cell cycle classifiers

An important means to test the performance of ccAFv2 is to compare it to existing state-of-the-art methods for cell cycle state classification. We evaluated the following methods: ccAF (O’Connor et al. 2021), ccSeurat (Hao et al. 2021), tricycle (Zheng et al. 2022), Revelio/SchwabeCC (Schwabe et al. 2020), reCAT (Liu et al. 2017), peco (Hsiao et al. 2020), and cyclone (Scialdone et al. 2015). We also evaluated the incorporation of ccAFv2 marker genes into the ccSeurat classification algorithm. However, it had significantly reduced performance compared to ccSeurat and ccAFv2 (**Supplemental Figure S2**). Each tool predicts a different subset of cell cycle phases, uses a different classification algorithm, was trained on different data, and requires different input genes and data formats (**Supplemental Table S4**). We applied ccAFv2 alongside the other state-of-the-art cell cycle classification methods to *in vivo* hNSCs collected from whole human fetal brain at PCW 9 R1 (**Figure 2A-B**; Zeng et al. 2023). These cells represent an independent dataset for an unbiased comparison of the cell cycle prediction algorithms. The hNSCs from Zeng et al., 2023 were also chosen for their similarity to the U5-hNSCs and their added real-world relevance, as they were collected *in vivo*. We chose to employ the ccSeurat classifier (Butler et al. 2018) to predict the reference labels for classifier comparison for the reasons described above. The AMI score is impacted by the number of cell cycle states in the reference (i.e., three cell cycle states in ccSeurat), and the number of states predicted by each algorithm (e.g., seven cell cycle states in ccAFv2). We used simulation studies to define the expected range of AMI scores that correspond to specific levels of similarity to the reference given the number of cell cycle states in the reference and the classifier being tested. The highest AMI was observed for tricycle, showing an 80% similarity to the reference (**Figure 2A**). This result aligns with the UMAP colorization, indicating a strong match within classifiers that predicted a comparable number of classes to ccSeurat (**Figure 2B**). reCAT and ccAFv2, predicting six and seven cell cycle states, respectively, achieved the next highest AMI scores, both demonstrating over 70% similarity to the reference (**Figure 2A**).

**Figure 2.**
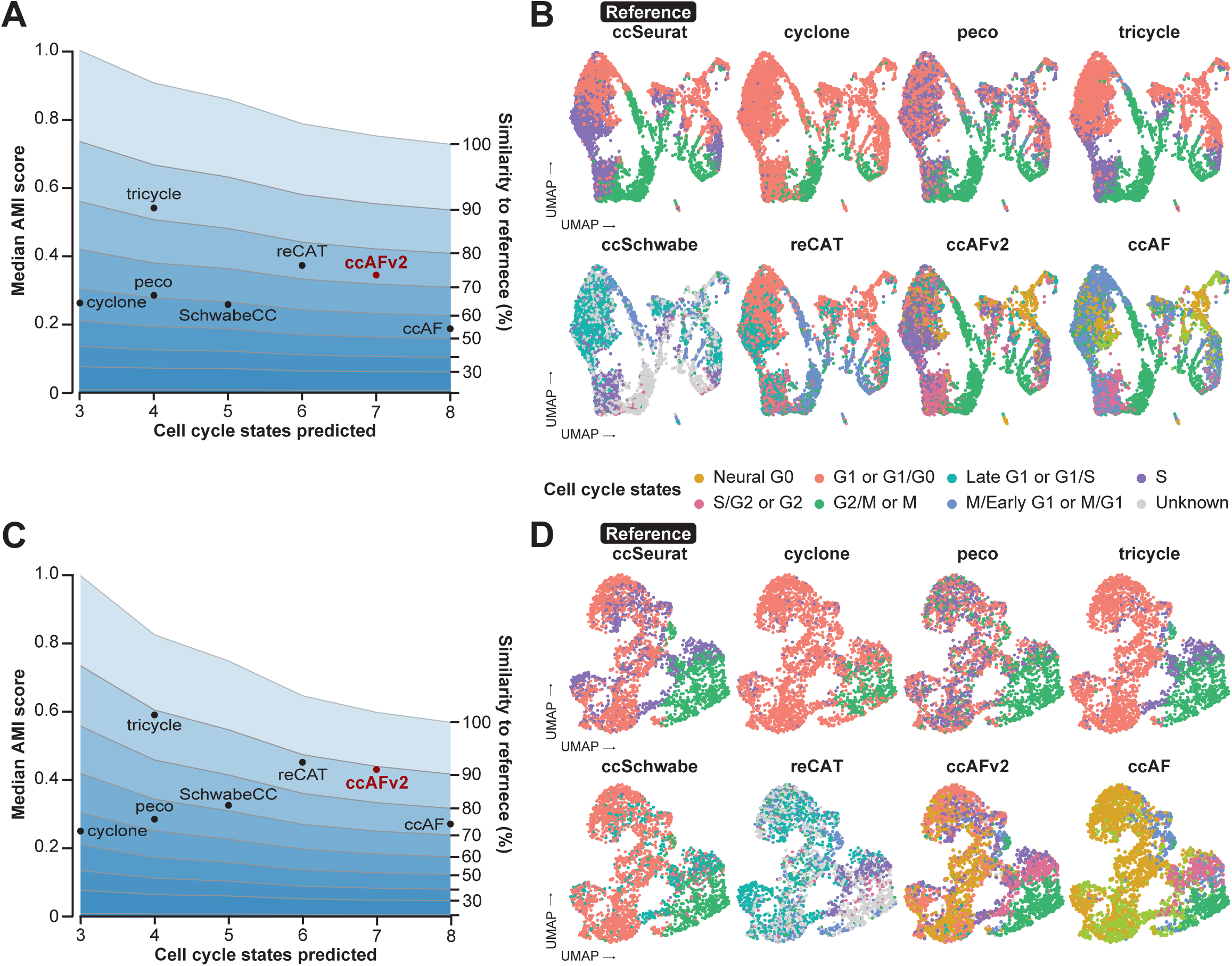
Comparing the performance of ccAFv2 to existing cell cycle state classifiers. **A.** Median AMI score for each cell cycle classifier’s predictions of the hNSCs from a whole fetal brain at 9 weeks post conception (PCW 9 R1; Zeng et al., 2023) relative to the ccSeurat cell cycle states is plotted against the number of cell cycle states predicted by the classifier. The average similarity to the reference was computed, based on the number of cell cycle states in the reference and predicted by the classifier, and were plotted at 10 percent intervals to facilitate comparison between classifiers with differing numbers of predicted cell cycle states. **B.** Overlay of representative cell cycle state predictions on the hNSCs from a whole fetal brain at PCW 9 R1. **C.** Median AMI score for each cell cycle classifier’s predictions of the glioma stem cell line BT322 relative to the ccSeurat cell cycle states is plotted against the number of cell cycle states predicted by the classifier. Again, average similarity to the reference was computed based on the number of cell cycle states in the reference and predicted by the classifier and were plotted at 10 percent intervals to facilitate comparison between classifiers with differing numbers of predicted cell cycle states. **D.** Overlay of representative cell cycle state predictions on the tumor cells of BT322.

Notably, ccAFv2 identified an S/G2 cluster of cells positioned between the S and G2/M cells classified by ccSeurat and tricycle, which is biologically plausible (**Figure 2B**). Additionally, while Neural G0 cells are intermixed with G1 and Late G1 cells within the proliferating cell population on the left side of the UMAP, the right side reveals a distinct cluster of Neural G0 cells (**Figure 2B**). This suggests the presence of a quiescent population in these normal human neural stem cells that is not detectable by the ccSeurat, tricycle, or reCAT classifiers.

We also applied ccAFv2 alongside the other cell cycle classification methods to cells derived from a glioblastoma (GBM) patient tumor (BT322; Couturier et al. 2020; **Figure 2C-D**). GBM patient tumors are characterized by both quiescent and proliferating subpopulations (Tejero et al. 2019) making them ideal datasets for evaluating and comparing different cell cycle classification methods. We used the ccSeurat labels as the reference because true cell cycle state labels do not exist for this dataset. Like the *in vivo* PCW 9 R1 hNSCs, the largest AMIs were observed for tricycle, reCAT, and ccAFv2; all of which correspond to just below 90% similarity to the reference (**Figure 2C**). These results demonstrate that ccAFv2 delivers at least equivalent performance when compared to contemporary state-of-the-art cell cycle classifiers, while providing the highest resolution of cell cycle state predictions including a quiescent-like G0 state (**Figure 2D**).

### *In vivo* cyclin expression and marker genes validate ccAFv2 cell cycle states

We explored the distribution of cell cycle states in 94,297 hNSCs collected from human fetal tissue at 3-12 weeks post-conception (Zeng et al. 2023; **Figure 3**). Application of ccAFv2 to the *in vivo* fetal hNSCs was found to differ by week stage (**Figure 3B**). The amount of Neural G0 cells from the *in vitro* U5-hNSCs, derived from fetal brain tissue at 8 weeks post-conception, (**Figure 3A**) matches closely to the *in vivo* hNSCs at eight weeks post-conception (**Figure 3B**). Moreover, the expression patterns of cyclins between the *in vitro* (O’Connor et al, 2021) and 5,575 *in vivo* hNSCs from PCW 9 R1 were similar (**Figure 3C-G**). In both hNSC populations, *CCNE2* exhibited its peak expression during the S phase, while *CCNA2* showed highest expression levels during the S/G2 and G2/M phases, and *CCNB1* displayed elevated expression in G2/M phase cells (**Figure 3G**). Notably, the highest expression of the key regulator of cell cycle progression, *CCND1*, was observed in the Late G1 state (**Figure 3G**).

**Figure 3.**
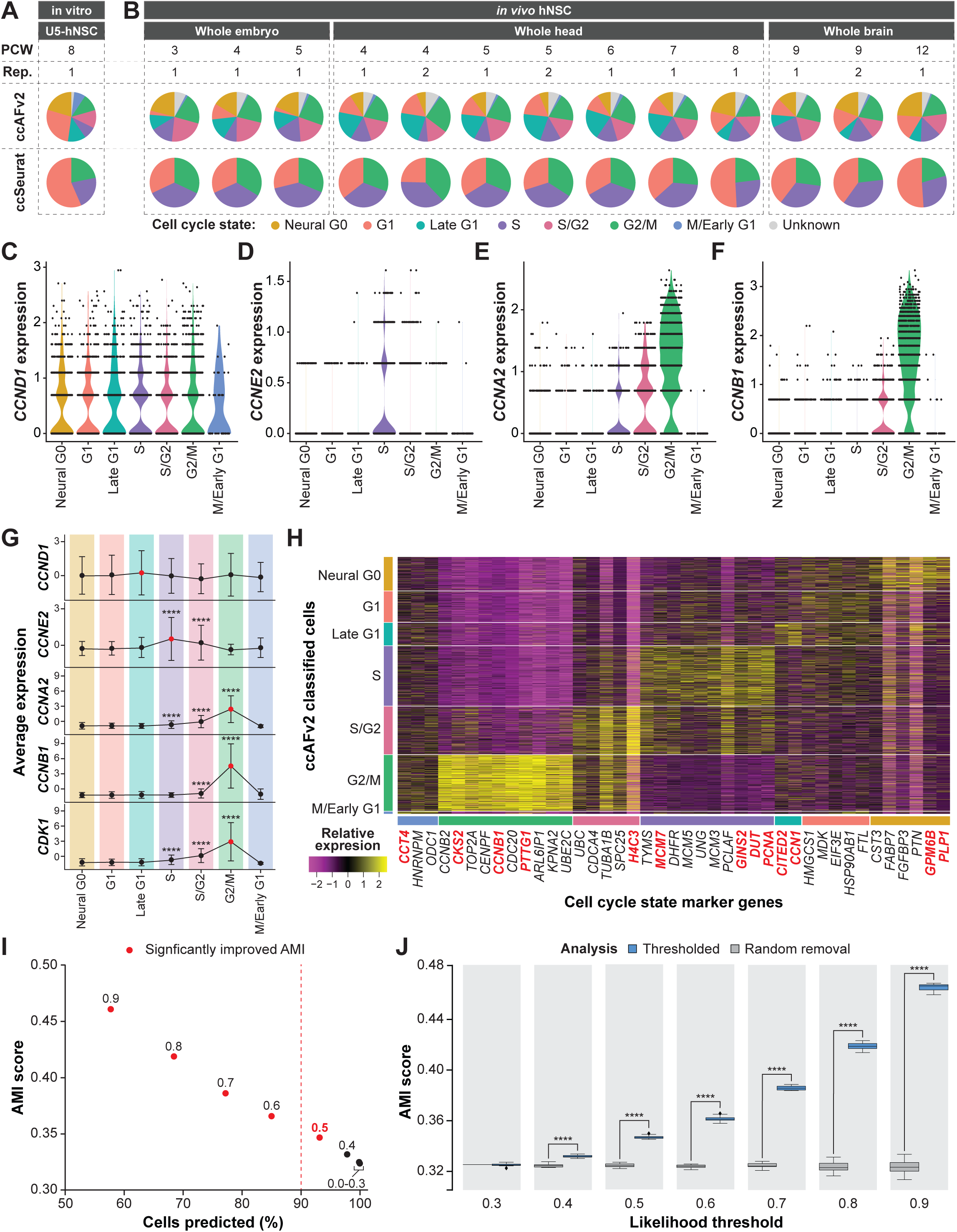
Application of ccAFv2 to *in vivo* hNSCs from fetal tissue 3 to 12 weeks post conception. **A.** Proportions of cell cycle states in U5-hNSCs which were grown *in vitro* and were derived from a human fetus at 8 PCW for both ccAFv2 and ccSeurat. **B.** Proportions of cell cycle states of hNSCs extracted from 3 to 12 PCW fetal tissue for both ccAFv2 and ccSeurat (Zeng et al, 2023). **C-F.** Distribution of cyclin expression in the *in vivo* hNSCs from a whole human fetal brain at PCW 9 R1 grouped by cell cycle phase. **G.** Mean expression of cyclins across the ccAFv2 cell cycle phases in cells from a whole human fetal brain at PCW 9 R1. Red points denote the ccAFv2 cell cycle state with the highest average expression. Gene expression levels at each cell cycle state were compared to those in G1 cells using Student’s *t*-test (**** indicates p ≤ 0.0001). **H.** Expression of ccAFv2 marker genes for each cell cycle state in hNSCs from a human whole fetal brain at PCW 9 R1. Important genes names are denoted in dark red. **I.** Testing different likelihood thresholds 0.0 to 0.9 using AMI score and percent of cells predicted as the metrics. Dashed red line indicates 90 percent of cells were predicted, and red dot indicates significantly improved AMI score due to applying threshold. **J.** Comparison of likelihood threshold application to random removal of the same number of cell predictions for *in vivo* hNSCs from a human whole fetal brain at PCW 9 R1. Metric used for assessment is the AMI score. Likelihood thresholds start at 0.3 on the x-axis because AMI values at likelihood thresholds 0 to 0.3 are the same. AMI scores at each likelihood threshold were compared using Student’s *t*-test (**** indicates p ≤ 0.0001). Rep. = biological replicate.

Additionally, we identified ccAFv2 marker genes that corresponded to cell cycle state markers in the PCW 9 R1 hNSCs. Differentially expressed genes for each cell cycle state were identified, and only those overlapping with the ccAFv2 marker gene lists were reported (**Figure 3H**).

Genes important to the ccAFv2 classifier were enriched among the translatable marker genes for PCW 9 R1 (**Figure 3H**). The expression patterns of these translatable marker genes were consistent with those observed in O’Connor et al., 2021. The exclusive or semi-exclusive expression of these markers in adjacent cell cycle states strongly supports the presence of high-resolution ccAFv2 clusters in hNSCs *in vivo*. Furthermore, the biological function of the translatable marker genes for each ccAFv2 cell cycle state validates the biological basis of the ccAFv2 clusters, providing further evidence of their relevance.

### Defining an appropriate classification likelihood threshold

The improved ccAFv2 classifier calculates likelihoods for each cell cycle state which can be used to determine the most likely state and to assess the quality of the classification. We hypothesized that applying a likelihood threshold to ccAFv2 classifications would ensure reliability and confidence in predicted cell cycle states by setting classifications for cells with less certainty to an “Unknown” state. We explored the range of possible likelihood thresholds on the 94,297 hNSCs collected by Zeng et al., 2023.

We tested ccAFv2 likelihood thresholds ranging from 0.0 to 0.9 in increments of 0.1 (**Supplemental Figure S3**). The calculated cell cycle state likelihood was required to be greater than or equal to the threshold, otherwise an “Unknown” state was returned (**Figure 1B)**. Each likelihood threshold was assessed using the percentage of cells predicted and an AMI score with ccSeurat cell cycle states as a reference. As the likelihood threshold increases the number of cells predicted decreases and the AMI scores increase (**Figure 3I**; **Supplemental Figure S3**; **Supplemental Table S5**). In other words, the removal of less certain classifications improves the accuracy of the overall classifications (**Figure 3I**). Next, we further demonstrated that the increase in AMI resulted from the specific removal of cells which had low classification likelihoods, by comparing it to the random removal of an equivalent number of cells (representative analysis for 9 weeks post-conception is shown in **Figure 3J**). The randomly removed cells do not increase the AMI (**Figure 3J**), only the selected removal of cells with low likelihoods were able to increase the AMI. We found that the median AMI scores calculated with likelihood thresholds of 0.4 to 0.9 were significantly higher than the median AMI scores of the randomly removed cells (**Figure 3J**; **Supplemental Table S5**), which indicates that the likelihood cutoffs of greater than or equal to 0.4 improve classification accuracy. We selected the likelihood threshold of greater than or equal to 0.5 because it signifies a minimum of 50% certainty in the classified cell cycle state. Additionally, greater than 90% of *in vivo* hNSCs could be assigned a cell cycle state with a likelihood threshold of 0.5 (**Figure 3I**). Thus, the threshold of 0.5 was set as the default for ccAFv2 and used in subsequent analyses, except where noted. We also provide users with the flexibility to adjust the likelihood threshold parameter in ccAFv2, allowing them to adapt the classifier’s operation to suit the unique characteristics of their dataset.

### Effect of missing gene expression values on ccAFv2

A known limitation of scRNA-seq is that dropouts are common. A dropout occurs when lowly to moderately expressed transcripts are detected in one cell but are not detected in another cell of the same cell type (Qiu 2020). Factors affecting dropouts include the number of sequencing reads from each cell and the complexity of the cell’s transcriptome. The ccAFv2 classifier uses the expression of 861 genes to predict cell cycle states. We hypothesized that dropouts could be simulated by randomly setting the expression of a defined percentage of genes to zero and that this would provide a reasonable approximation of the influence of missing genes on the accuracy of ccAFv2’s cell cycle state classifications. We evaluated the consequences of these simulated gene dropouts on the classifier error rate, AMI, and the number of cells predicted (**Supplemental Figure S4**-**5**; **Supplemental Table S6**-**7**). As described earlier, the median error rate of applying ccAFv2 to U5-hNSCs was 3.3% with 99% of the input genes (99% is used to allow for cross-validation). Missing information for 20% of the ccAFv2 input genes yielded a smaller median error rate (12.2%) than the original ccAF error rate with all the input genes (18.4%), underscoring the improved performance of the new model. Introducing missing information for 40% of ccAFv2 input genes led to a 29% median error rate, and 96% of cells were predicted (**Supplemental Table S6**). The error rate was the most affected by the introduction of missing information (**Supplemental Figure S4A)** and the median percentage of cells predicted remained above 80% even when 70% of the input gene list was set to missing (**Supplemental Table S6**). When breaking down the error rate by cell cycle state, we observed that S and M/Early G1 had the highest error rates as missing information increased (**Supplemental Figure S4B**; **Supplemental Table S7**). However, the number of cells predicted remained relatively consistent across all states despite the increasing in missing data (**Supplemental Figure S4C**; **Supplemental Table S7**). The increase in error rate without a concomitant decrease in the number of cells predicted suggests that raising ccAFv2’s likelihood threshold (>0.5) might be required to ensure the quality of predictions for datasets with greater than 20% missing ccAFv2 input genes. Indeed, the error rate for introducing 20% missing information decreased from 12.2% median error rate at 0.5 likelihood threshold to 9.9% with a 0.7 likelihood threshold (**Supplemental Figure S4D**; **Supplemental Table S6**) and 6.2% with a 0.9 likelihood threshold (**Supplemental Figure S4E**; **Supplemental Table S6**). Thus, introducing 20% missing information led to four times the error rate, and the increased error rate can be mitigated in part by increasing the likelihood threshold. Increasing the likelihood threshold decreases the error rate by removing classifications for cells where the missing information has degraded the confidence in the prediction. By removing predictions with less confidence, the error rate decreases, but the overall number of cells classified with cell cycle states decreases. Testing the impact of increasing the likelihood threshold on the number of predicted cells can be quite insightful for choosing an appropriate likelihood threshold (**Supplemental Figure S4F**). Careful consideration of the balance between minimizing errors and retaining enough cells for downstream studies is essential.

### Neural G0 state is enriched in mesenchymal G0 cells

We developed an experimental method to isolate fixed G0 cells using fluorescence-activated cell sorting (FACS) with established markers (Gookin et al. 2017). This approach ensures the selected cells are diploid, non-replicating, and have unphosphorylated RB (pRB^-^), all while preserving RNA integrity. The experimental approach was applied to identify G0 cells from human skeletal muscle satellite cells (hSkMSCs), which are derived from the mesodermal germ layer (**Figure 4A**). Subsequent RNA-seq of the sorted G0 cells captures the characteristic expression pattern of the G0 state for that cell type, enabling direct comparison with the expression profiles of single cells from scRNA-seq data of unsorted cells. We characterized the bulk RNA-seq signatures of 400,000 G0 hSkMSCs for two biological replicates. These G0 signatures were mapped onto 6,921 asynchronous, unstained, and unsorted hSkMSCs collected using scRNA-seq. Next, the ccAFv2 cell cycle classifier was applied to the hSkMSC scRNA-seq data. Cells in S, S/G2, G2/M and M/Early G1 states formed distinct clusters ordered in the canonical cell cycle patterning, highlighting the generalizability of these ccAFv2 states to *in vitro* hSkMSCs (**Figure 4B**). The Late G1 cells did form a cluster that was positioned in front of the S phase cells, however, additional Late G1 cells can be seen dispersed within the left half of the cells. This dispersion may be partly attributed to the higher *CCND1* expression in hSkMSCs G0 cells compared to hNSC Neural G0 cells (**Supplemental Figure S6**. The cells inside the dashed region contain almost exclusively Neural G0, G1, and Late G1 cells and do not coalesce into defined clusters (**Figure 4B-C**), suggesting that the ccAFv2 classifier was struggling to accurately discriminate between Neural G0, G1, and Late G1. Correlation of the experimentally determined G0 signature to the cells from the scRNA-seq revealed significant enrichment within the ccAFv2-labeled Neural G0 and G1 cells (**Figure 4D-F**). The majority of G0 cells were classified as Neural G0, while misclassified G0 cells were predominantly labeled G1 or Late G1, and very few were classified as the cycling states (S, S/G2, G2/M, M/Early G1; **Figure 4G**). These findings confirm that ccAFv2 is having difficulty discriminating between the Neural G0, G1, and Late G1 states in hSkMSCs derived from the mesoderm dermal layer.

**Figure 4.**
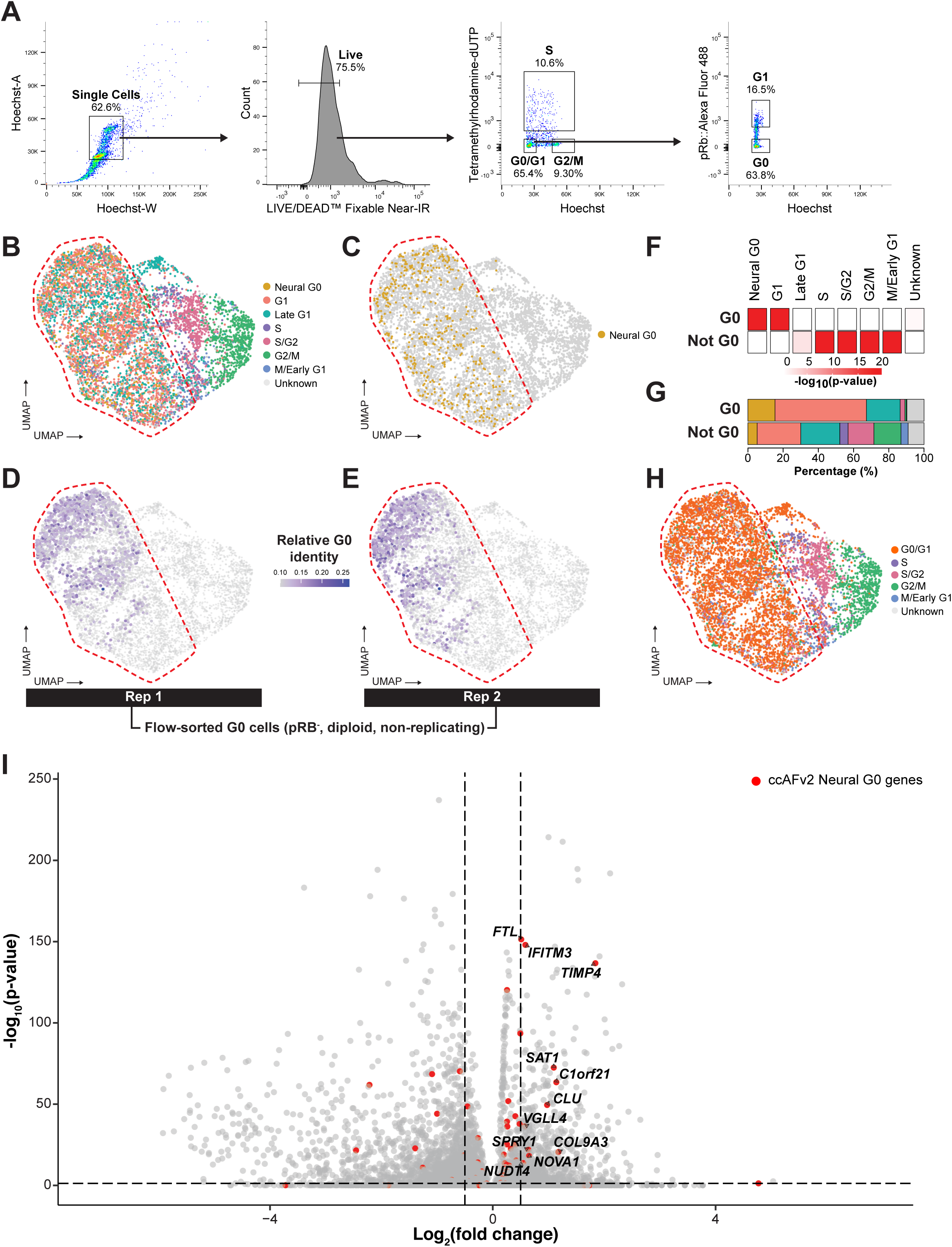
Experimental enrichment of mesenchymal G0 cells from hSkMSCs using FACs. **A**. Gating strategy for isolating mesenchymal G0 cells from hSkMSCs. First gated on single cells, then live cells, next diploid cells that are not replicating are selected, and finally cells with hypo-phosphorylation of RB are selected. **B**. ccAFv2 applied to unsorted hSkMSCs. The red dashed area encompasses the UMAP area containing the vast majority of Neural G0, G1, and Late G1 cells, these three states do not exhibit consistent clustering. **C**. Neural G0 cells are highlighted in color, while all other states are shown in gray. **D-E**. Cells are colored based on their Spearman correlation coefficient with the scRNA-seq expression profiles and the RNA-seq profiles of flow-sorted mesenchymal G0 cells (pRB-, diploid, non-replicating) from two biological replicates. **F**. Test of which ccAFv2 cell cycle states were significantly enriched with mesenchymal G0 cells, and not mesenchymal G0 cells. Mesenchymal G0 cells are defined by a Spearman correlation ≥ 0.1 in both replicates, while non-G0 cells are defined by a correlation ≤ 0.1 in one or both replicates. Values are represented as the negative logarithm of the p-value. **G**. Percentage of ccAFv2 states in mesenchymal G0 and not G0 cells. **H**. Differential expression of genes between mesenchymal G0 cells versus not G0 cells. Each dot represents one gene (n = 22,845). Dotted lines denote log2(fold change) and adjusted p-value cutoffs to identify significant marker genes (log2FC ≥ 0.5; p-adj ≤ 0.05). Red dots denote ccAFv2 Neural G0 marker genes. Labeled genes are marker genes for hSkMSC G0 cells that overlap with ccAFv2 Neural G0 marker genes.

However, the misclassifications are systematic rather than random, and a straightforward solution of merging Neural G0, G1, and Late G1 classifications effectively resolves the misclassifications. To accommodate this, we implemented a switch in the ccAFv2 classifier, enabling users to choose whether to combine Neural G0, G1, and Late G1 or to keep them separate. This feature provides a more cautious and flexible approach for classifying cell types beyond neuroepithelial cells (**Figure 4H**). This provides users with the flexibility to use ccAFv2 higher resolution cell cycle classification for non-neuroepithelial cell types.

Additionally, it should be noted that while the Neural G0 state does not accurately capture the G0 state of hSkMSCs the experimental data demonstrates that it is possible to identify a subpopulation of G0 cells. The marker genes discovered for the hSkMSC G0 cells were significantly overlapping with the Neural G0 marker genes (n = 11; p-value = 4.4 x 10^-4^; *FTL*, *IFITM3*, *TIMP4*, *SAT1*, *C1orf21*, *CLU*, *VGLL4*, *SPRY1*, *COL9A3*, *NOVA1*, *NUDTA*; **Figure 4I**; **Supplemental Table S8**. Which strongly suggests that it may be possible to train a classifier with a more generalizable G0 state in future studies using our experimental approach to characterize new training datasets.

### ccAFv2 cell cycle states are generalizable across germ layers

Another key consideration when using ccAFv2 is its ability to accurately predict cell cycle states (S, S/G2, G2/M, and M/Early G1) in cell types beyond neuroepithelial cells. In Zeng et al., 2023, they profiled single cells from human fetal tissues, representing all three germ layers (endoderm, mesoderm, and ectoderm; **Figure 5A**). We applied ccAFv2 to 245,906 cells from the atlas first including Neural G0, G1 and Late G1 predictions (**Figure 5B**), and then by collapsing these three predictions into a G0/G1 class (**Figure 5C**). These representations of the Zeng et al., 2023 dataset aggregate across time (PCW 3–12) and tissue collection strategies (whole embryo, whole head, and brain). The proportions of cell cycle states across time for each cell type show strong concordance (**Supplemental Figure S7**), highlighting the consistency of ccAFv2 predictions across biologically similar independent scRNA-seq datasets. Next, we analyzed cyclin expression patterns across the 15 distinct cell types and calculated the average expression for each germ layer. Each germ layer exhibited the expected cyclin expression pattern, with *CCND1* driving entry into the cell cycle (**Figure 5D**), *CCNE2* peaking during S phase (**Figure 5E**), and *CCNA2* (**Figure 5F**) and *CCNB1* coordinating progression and regulation of cell division (**Figure 5G**). These results provide strong evidence that the ccAFv2 predicted S, S/G2, G2/M, and M/Early G1 states are accurate across cell types derived from all germ layers.

**Figure 5.**
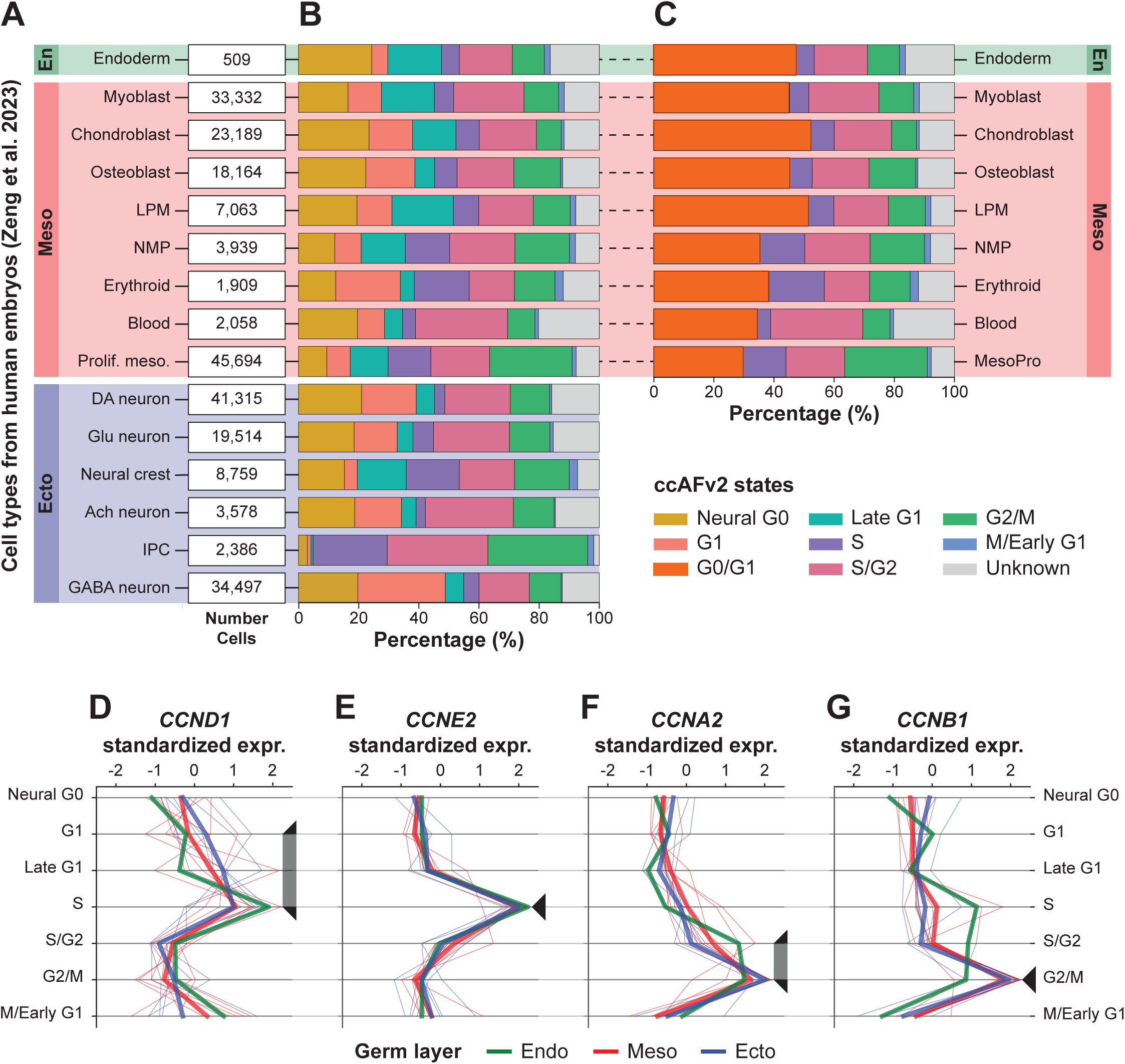
Application of ccAFv2 to the transcriptomes of 245,906 single cells derived from human fetuses aged 3 to 12 PCW. **A**. The 15 different cell types included in the analysis encompass all three germ layers. For each cell type the number of cells is given. **B**. Percentage of each ccAFv2 predicted state for each cell type. **C**. Percentage of each ccAFv2 predicted state for each cell type when Neural G0, G1, and Late G1 are binned. **D-G**. Z-score normalized cyclin expression across 15 cell types. Thin lines represent individual cell types, while thick lines indicate the average Z-score normalized cyclin expression for each germ layer. Lines are color-coded according to their corresponding germ layer.

We also observed that ccAFv2 proportions align closely with expected developmental patterns. For instance, during PCW 9–12, the fetal brain undergoes rapid development, marked by significant cell division as major structures like the cerebrum, cerebellum, and brainstem become more defined (Belmonte-Mateos and Pujades 2021; Martínez-Cerdeño et al. 2006). We found that intermediate progenitor cells (IPCs) detected at PCW 9 and 12 in whole brains were predominantly in the S, S/G2, and G2/M phases, consistent with the active proliferation necessary for forming these brain structures (**Figure 5**; **Supplemental Figure S7**).

Furthermore, these PCW 9–12 IPCs exhibited high expression of EOMES, a critical factor that drives the expansion of the IPC pool (Arnold et al. 2008). The much-reduced proportions of Neural G0, G1, and Late G1 in IPCs validates that ccAFv2 predictions are consistent with known biology. We also observed reduced Neural G0, G1, and Late G1 proportions in the non-neuroepithelial proliferating mesoderm (Prolif. meso.; **Figure 5**) defined by high expression of the proliferation marker *MKI67* (log2(FC) ≥ 1.24) and mesoderm marker *CDH11* (log2(FC) >2.27) (Hoffmann and Balling 1995). The reduced number of non-cycling cells in IPCs and proliferating mesoderm cell types is well documented and demonstrates that while Neural G0, G1 and Late G1 misclassifications may occur that the relative proportions of non-cycling cells to cycling cells is accurately determined by ccAFv2.

### Capturing the effect of growth factors on cellular proliferation

Growth factors are used to increase cellular proliferation *in vitro*, and we characterized the transcriptomes of LGG cells (grade 2 astrocytoma and grade 3 oligodendroglioma) with and without the application of growth factors (**Supplemental Figure S8**). For this analysis we tested the impact of adjusting the ccAFv2 likelihood threshold across a range of values 0 to 0.9 (**Supplemental Figure S9A**-**B**). Increasing the likelihood threshold values from 0.4 to 0.9 led to an increased proportion of “Unknown” classifications in the samples without growth factors, which is consistent with the known effect of growth factors to stimulate proliferation and the cell cycle. The increased proportion of “Unknown” cells may correspond to new growth factor starvation state(s) not included in ccAFv2 classification states. Additionally, the S, S/G2, and G2/M cell cycle states were disproportionately removed as the likelihood threshold increased (**Supplemental Figure S9A**-**B**). We then set the likelihood threshold to 0.9 and observed that the cells grown with growth factors form clusters of cell cycle state labels, outlining the expected progression of cell cycle phases (G1 → S → S/G2 → G2/M → M/Early G1; **Supplemental Figure S8A**, **C**, **D**, & **F**). Conversely, cells grown without growth factors exhibit a more dispersed distribution of cell cycle state labels (**Supplemental Figure S8B**, **C**, **E**, & **F**). The ability to change the likelihood threshold of ccAFv2 allows us to observe the biological impact of adding growth factors to LGG cells and demonstrates what to expect when the cell cycle is not the main transcriptional signal in cells.

### Removing cell cycle expression signatures

The cell cycle generates a strong transcriptional signature that can obscure other less robust transcriptional signatures of interest. Previous studies have shown that statistical methods can effectively remove cell cycle transcriptional signatures, and that the residual transcriptional variance can be used to study less robust transcriptional signatures of interest (Luecken and Theis 2019). We showcase successful removal of the cell cycle transcriptional signatures for the U5-hNSCs, and LGG cells. First, each cell cycle state’s average expression of marker genes is computed for every single cell or nuclei. Then, these average cell cycle expression patterns are regressed out of the dataset during normalization. The ccSeurat regression method uses only the S and G2/M cell cycle states, so we first tested regression with the S and G2/M cell cycle states from ccAFv2. We found that the ccAFv2 marker gene derived average cell cycle expression patterns could mitigate cell cycle transcriptional signatures as effectively as ccSeurat (empirical p-value > 0.05, **Supplemental Table S9**; **Supplemental Figure S10**). Additionally, we found that incorporating Late G1, S, S/G2, G2M, and M/Early G1 was also quite effective and led to a more robust homogenization of the cell cycle states based on PCA plots (**Supplemental Table S9**; **Supplemental Figure S10**). This approach enables researchers to dissect complex gene expression patterns and uncover novel insights into cellular processes beyond the cell cycle.

### Classifying neuroepithelial-derived cells in humans and mice

It was crucial for ccAFv2 to be highly user-friendly, ensuring researchers can easily apply it across a wide range of datasets. The model was designed to accept inputs for tissue source, data type, and gene identifier, eliminating the need for manual data conversion (**Figure 6A**). In a previous study (O’Connor et al. 2021) we applied the ccAF classifier to cells from the developing human telencephalon (Nowakowski et al. 2017). We applied ccAFv2 to these same cells and compared the ccAF and ccAFv2 predicted cell cycle proportions. We observed that the Neural G0 state was less frequent in all cell types for ccAFv2 relative to ccAF (**Figure 6B**; **Supplemental Table S10**). The Neural G0 state was distinctly less frequent in the neuronal cell types. For ccAF Neural G0 made up most of the cell cycle states for EN-PFC and EN-V1, but in ccAFv2 these two cell types classified primarily as G1 (**Figure 6B**). The glial cell types had the largest Neural G0 subpopulations (**Figure 6B**).

**Figure 6.**
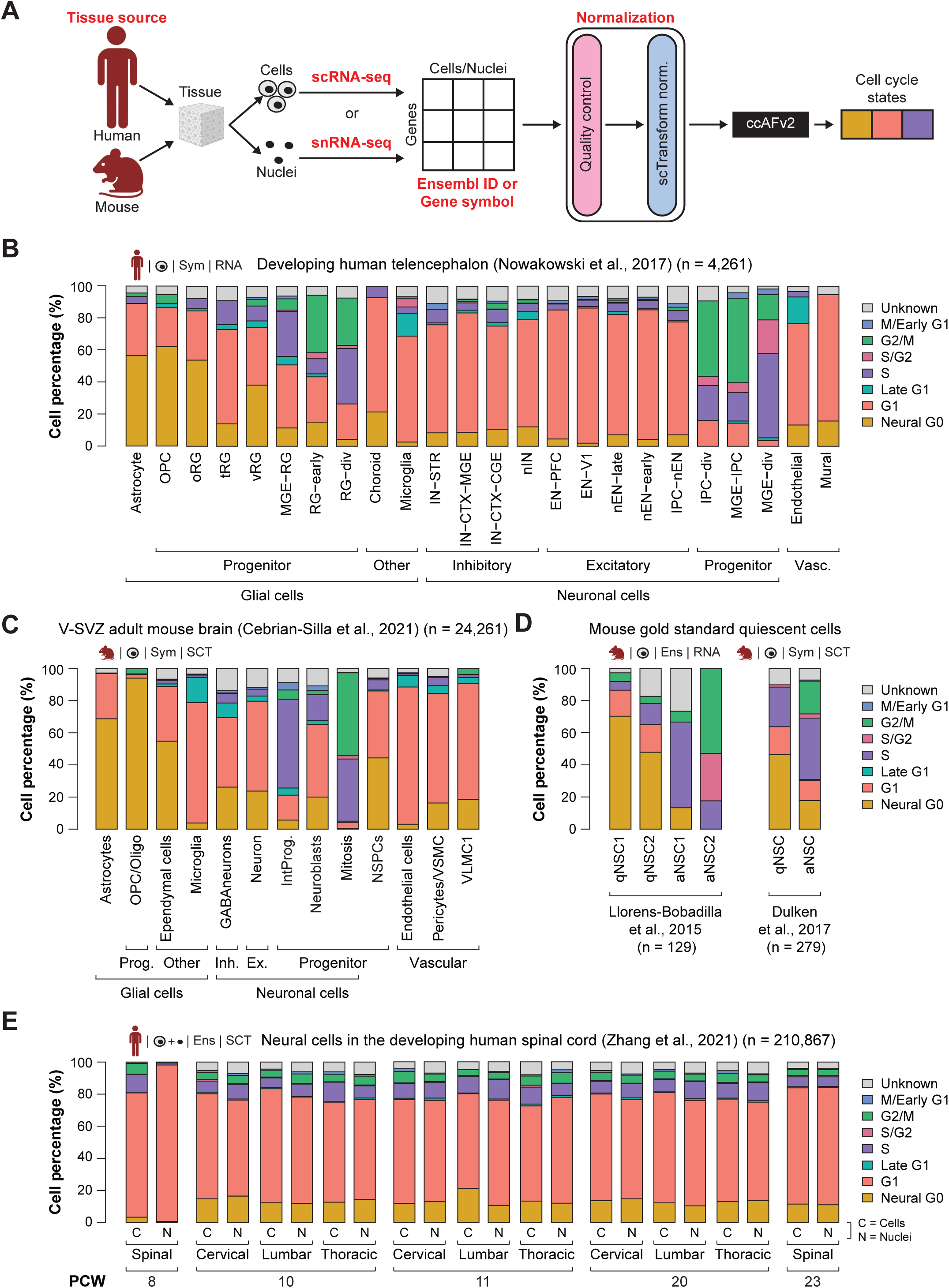
Application of ccAFv2 to single cells and nuclei from human and mice. **A.** Summary schematic of data ccAFv2 can be applied to and suggested data preparation. **B.** Proportion of cells assigned to each cell cycle state for scRNA-seq data from the developing human telencephalon (Nowakowski et al, 2017). **C.** Proportions of cell cycle states from scRNA-seq from the ventricular (V)-SVZ of the adult mouse brain (Cebrian-Silla et al, 2021). Prog. = progenitors, Inh = inhibitor, Ex = excitatory, NSPCs = neural stem/progenitor cells, IntProg. = intermediate progenitor cells. **D.** Proportions of cell cycle states from scRNA-seq from GLAST and PROM1 flow-sorted cells from the subventricular zone (SVZ) of mice (Llorens-Bobadilla et al, 2015), and EGFR, GFAP, and PROM1 flow-sorted cells from the subventricular zone (SVZ) of adult mice (Dulken et al, 2017). qNSC1 = dormant quiescent neural stem cell, qNSC2 = primed-quiescent neural stem cell, aNSC1 = active neural stem cell, aNSC2 = actively dividing neural stem cell. qNSC = quiescent neural stem cell, aNSC = active neural stem cell. **E.** Proportions of cell cycle states from scRNA-seq (C) and snRNA-seq (N) from spinal, cervical, lumbar, and thoracic regions from the developing human spinal cord at 8, 10, 11, 20, and 23 PCW (Zhang et al, 2021).

We also applied ccAFv2 to cells from the ventricular-subventricular zone (V-SVZ) of the adult mouse brain (Cebrian-Silla et al. 2021), a location known to contain neural stem and precursor cells in the adult brain (Lim and Alvarez-Buylla 2016). The adult mouse V-SVZ validates observations from the developing human telencephalon (**Figure 6C**). In the V-SVZ the glial cells tended to have larger Neural G0 subpopulations, neuronal cell types tended to have less Neural G0, and microglial had the smallest amount Neural G0 (**Figure 6C**). The results are similar given the differences between species, developmental state, and anatomical origins. These findings illustrate that the ccAFv2 classifier can be applied to cells originating from both humans and mice.

### Classifying quiescent-like neural stem cells

Previously we validated the Neural G0 state using two independent *in vivo* scRNA-seq profiling studies of NSCs from adult neurogenesis in the subventricular zone (SVZ) that used fluorescence activated cell sorting (FACS) to sort out quiescent and activated NSCs (Llorens-Bobadilla et al. 2015; Dulken et al. 2017). We applied ccAFv2 to these same cells and compared the ccAF and ccAFv2 predicted Neural G0 subpopulations. Overall, the qNSCs are enriched with quiescent-like Neural G0 cells, and the aNSCs are at some stage of the cell cycle (**Figure 6D**). The proportion of cells classified as Neural G0 decreased for ccAFv2 in the quiescent NSCs (qNSCs) and was replaced by more G1, S/G2, and a small amount of G2/M (**Figure 6D**; **Supplemental Figure S11**). For Llorens-Bobadilla et al., 2015 the active NSCs 1 (aNSC1) were more highly enriched with S phase cells, and aNSC2 were enriched with S/G2 and G2/M. A similar trend was observed for the Dulken et al., 2017 dataset. Additionally, we used the G0 arrest signature from Wiecek et al., 2023 to validate the Neural G0 state in ccAFv2 (Wiecek et al. 2023). We found significant enrichment of the G0 arrest signature (i.e., QuieScore G0) within the U5-hNSC Neural G0 and G1 states (**Supplemental Figure S12**). These results continue to validate our assertion that Neural G0 represents a quiescent-like cell state, and that ccAFv2 can accurately classify this quiescent-like G0 state.

### Accurate classification of cells and nuclei

Tissues in single-cell studies can be processed into cells for scRNA-seq or nuclei for snRNA-seq. Both methods are commonly used and have advantages and limitations (Slyper et al. 2020). Thus, it is important to demonstrate whether ccAFv2, which is trained on cells, can accurately classify cell cycle states for single nuclei. We employed the Zhang et al., 2021 dataset which characterized developing human spinal cord tissue from five developmental time points using both scRNA-seq and snRNA-seq from the same experimental conditions (Zhang et al. 2021). The proportions of cells in each cell cycle state are similar between scRNA-seq and snRNA-seq from the same condition (**Figure 6E**), illustrating the versatility of the ccAFv2 classifier in effectively analyzing both scRNA-seq and snRNA-seq profiles.

### Mapping proliferation onto tissue through spatial transcriptomics

scRNA-seq and snRNA-seq provide valuable information about the transcriptional states of cells and nuclei, but without contextual information, relating these states with previously described biology can be challenging. Spatial transcriptomics captures the transcriptional activity of a single-cell or a region containing a small number of cells at a position within an intact tissue, offering structural information that can be referenced to anatomical atlases and established histology. We applied ccAFv2 to the highest resolution sequencing-based spatial transcriptomic dataset currently available, derived from a slice of a mouse E15.5 embryo binned to 8µm, achieving near cellular resolution (**Figure 7A-B**). Despite rapid development at late prenatal stages, Neural G0 classified spots were highly prominent, particularly across the midbrain region. The proliferation marker *Mki67* was minimally expressed in these G0 enriched areas and, in the brain, highlighted the described stem cell niches of the lower, medial, and caudal ganglionic eminences (LGE, MGE, CGE), along with the stem cell migratory paths from these regions along the sub-ventricular zone (SVZ; **Figure 7C**; Kriegstein and Noctor 2004).

**Figure 7.**
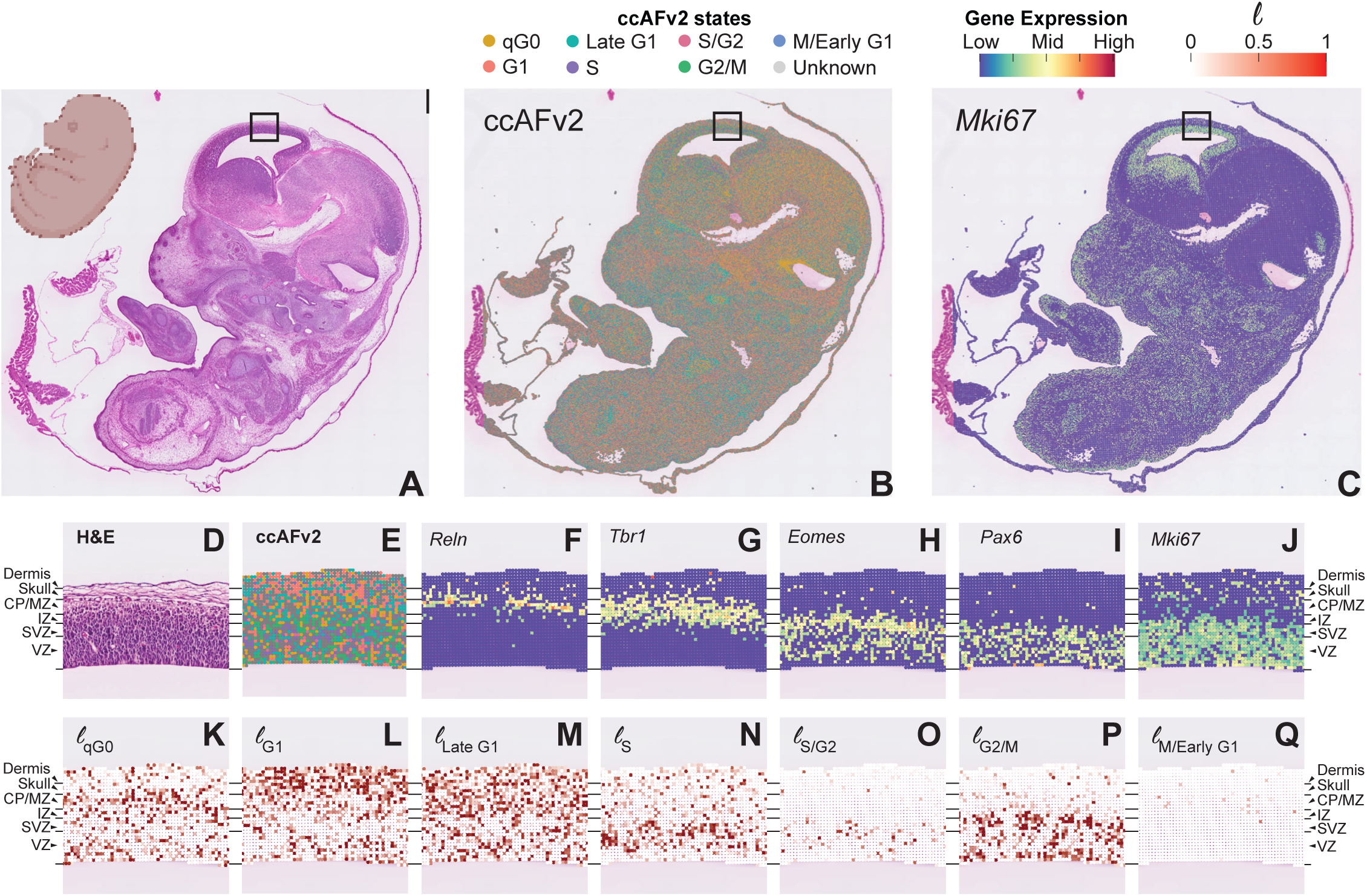
Application of ccAFv2 to spatial transcriptomics data from a male C57BL/6 mouse embryo at E15.5. **A.** H&E staining for the whole embryo. **B**. Spatial overlay of the predicted ccAFv2 states onto the whole embryo. **C.** Spatial expression of the cell cycle marker gene *Mki67* for the whole embryo. The black boxes in panels **A** through **C** indicate the region of the developing cortex that was magnified in panels **D** through **Q**. The developmental regions of the developing cortex are denoted on the side: Dermis = developing skin, Skull = developing skull, CP/MZ = cortical plate and marginal zone, IZ = intermediate zone, SVZ = subventricular zone, VZ = ventricular zone. **D.** H&E staining for the developing embryo cortex. **E.** Spatial overlay of the predicted ccAFv2 states onto the developing embryo cortex. **F**-**I.** Expression of key marker genes describing the developmental regions in the developing embryo cortex. **J.** Spatial expression of the cell cycle marker gene *Mki67* in the developing embryo cortex. **K**-**Q.** Likelihoods for each of the cell cycle states spatially overlayed onto the developing embryo cortex. The magnitude of the likelihood indicates the probability that a cell with that cell cycle state underlies that spot of the spatial array.

From E12.5 to E17.5 the mouse cortex develops in a well-defined layers (**Figure 7D**). Applying ccAFv2 to the cortex captured the layered patterning of cell cycle states that fit with our current model of cortical development (**Figure 7E-Q**). Histology showed an outer layer of dermis (skin) marked by *Krt5* expression (**Supplemental Figure S13A**-**B**), covering the developing skull identified by *Col1a1* expression (**Supplemental Figure S13C**), followed by densely packed cells of the brain (**Figure 7D**). Precursors of excitatory neurons migrate along and divide in the SVZ with asymmetric division specifically occurring within the ventricular zone (VZ). A much smaller population of inhibitory neuronal precursors migrate and divide along the intermediate zone (IZ) and medial zone (MZ) before invading the cortical plate (CP) and differentiating into their neuronal sub-type. By E15.5, the CP is already well populated with post-mitotic neurons that previously migrated from the SVZ from E12.5-E14.5 and will form layers IV-VI of the adult cortex. Upon reaching the border of the CP and MZ, neural stem cells receive maturation factors from glial cell types including *Reln* (**Figure 7F**), with canonically post-mitotic neurons marked by *Tbr1* (**Figure 7G**; Englund et al. 2005). This post-mitotic region classifies as Neural G0 by ccAFv2 (**Figure 7E** & **K**). Similarly, *Mki67* is sparse within the CP, but highly active in the SVZ and VZ (**Figure 7J**). Intermediate progenitor cells (IPCs) in the SVZ, marked by *Eomes*, undergo symmetric division before radiating outward (**Figure 7G**). Within the VZ radial glia migrate further inward before asymmetric division, with newly divided IPCs radiating back up to the SVZ and outward to populate the cortex. These events are captured by ccAFv2 as a single enriched S and S/G2 band marking the SVZ (**Figure 7N** & **O**). We also observed two bands containing G2/M classified spots, corresponding to regions of symmetric division in the SVZ and asymmetric division in the VZ (**Figure 7P**). Cells committed to differentiation, immediately migrate outward along and the few spots that classify as M/Early G1 were almost entirely in the IZ and above (**Figure 7Q**). These results demonstrate that ccAFv2 can be effectively applied to spatial transcriptomics of the developing cortex, accurately recapitulating known biological insights into the spatial organization of proliferative activity.

## Discussion

We designed ccAFv2 to use transcriptomic data to accurately classify cell cycle states and a quiescent-like G0 state for single cells or nuclei. The performance of the updated classifier was superior to its predecessor and demonstrated comparable or better performance than other state-of-the-art cell cycle classifiers. The ccAFv2 classifies cells into a broader range of cell cycle states than the contemporary state-of-the-art cell cycle classifiers (Hao et al. 2021; Zheng et al. 2022; Schwabe et al. 2020; Liu et al. 2017; Hsiao et al. 2020; Scialdone et al. 2015) and includes a quiescent-like G0 state. Moreover, ccAFv2 features a tunable parameter to filter out less certain classifications. We showcased the versatility of ccAFv2 by successfully applying it to classify cells, nuclei, and spatial transcriptomics data in humans and mice, using various normalization methods and gene identifiers. The classifier can be used either as an R package integrated with Seurat (https://github.com/plaisier-lab/ccafv2_R) or a PyPI package integrated with SCANPY (https://pypi.org/project/ccAF/). We proved that ccAFv2 has enhanced accuracy, flexibility, and adaptability across various experimental conditions, establishing ccAFv2 as a powerful tool for exploring cell cycle dynamics in diverse biological contexts.

A major limitation of developing cell cycle classifiers is a lack of scRNA-seq datasets with ground truth labels for each cell cycle state, including G0. Previous studies have used the DNA stain Hoechst (Buettner et al. 2015) or FUCCI (Leng et al. 2015) to sort cells into G1, S, and G2M subpopulations. However, the limitations of these studies render them unsuitable for constructing a classifier. These limitations include the use of embryonic stem cells which lack distinct G1 or G0 phases as the model system (Ballabeni et al. 2011; White and Dalton 2005), not of fixing the cells, and reliance on non-transcriptional markers. We demonstrated the ability to isolate cells from specific cell cycle states, characterize them transcriptionally, and map their signatures onto scRNA-seq data to identify the spatial distribution of those states. In future studies, we aim to leverage this approach to create training datasets for developing more robust and generalizable cell cycle classifiers.

Batch effects pose a significant challenge for single cell, nuclei, and spatial RNA-seq studies; and we have addressed their impact on ccAFv2 in three ways. First, we strongly recommend users apply ccAFv2 to each dataset separately prior to any integration or combining of datasets. The ccAFv2 predictions in the metadata integrate very easily and preempt any issues caused by batch effects between datasets. Second, we advise using SCTransform normalization to standardize datasets to the Pearson residual scale, which mitigates differences in magnitude and variance (Hafemeister and Satija 2019). The ccAFv2 R package is specifically parameterized to reapply SCTransform, normalizing the expression of all genes, not just the most variable ones, to maximize overlap with ccAFv2 marker genes, ensuring optimal classification accuracy for each cell. Finally, ccAFv2 model incorporates expression of multiple marker genes to predict a cell cycle state. This approach minimizes the influence of batch effects, as a significant misclassification would require the simultaneous, directional alteration of multiple marker genes, a highly improbable scenario for random batch effects. By adhering to these recommendations and leveraging the design of ccAFv2, we provide effective strategies to mitigate the impact of batch effects when using ccAFv2.

The ccAFv2 classifier will be most helpful in biological contexts where the cell cycle is active. We utilized atlases of developing human and mouse embryos and fetuses because proliferation is essential in developing organisms (Soufi and Dalton 2016; Pauklin and Vallier 2013).

Evidence is building to show that cell fate decisions are tightly coupled to cell cycle events and machinery (Pauklin and Vallier 2013). In healthy adult organisms, proliferation plays critical roles in several processes: maintenance of stem cell populations (Harada et al. 2021), clonal expansion of both innate and adaptive immune cells (Adams et al. 2020), and germ cell meiosis, encompassing oogenesis (Bukovsky et al. 2005) and spermatogenesis (Guo et al. 2018). Our recommendation is to aggregate Neural G0, G1, and Late G1 into G0/G1 when applying ccAFv2 to cell types that are not neuroepithelial. Defects in cell cycle machinery or regulation can lead to runaway proliferation characteristic of cancer (Hanahan and Weinberg 2000), or the lack of proliferation of crucial cell types can lead to neurodegenerative disorders (Joseph et al. 2020).

Cell cycle classification would benefit any *in vitro*, *in vivo*, or *ex vivo* studies of proliferating cells. On the other hand, we provide methods to regress the cell cycle expression patterns out of single cell or nuclei data to uncover underlying biological signals. Overall, incorporating cell cycle states into single-cell and nuclei studies enhances our ability to dissect complex biological systems, unravel cellular heterogeneity, and decipher the molecular mechanisms by which proliferation affects cellular processes.

The studies reported here demonstrate that the quiescent-like G0 state (Neural G0) in ccAFv2 is detectable across all three germ layers in developing human fetal cells and provided a list of putative marker genes that are common across cell types. This corroborates our previous findings that Neural G0 was an active transcriptional signature executed by a subpopulation of U5-hNSCs (O’Connor et al. 2021). However, support for novel G0 states was observed in the growth factor deprived LGG cells, where an increased proportion of “Unknown” cells was detected, hinting at novel quiescent-like state(s) missing from ccAFv2. Other studies have identified multiple G0 states in a single cell type that are invoked in response to different stimuli (e.g., spontaneous loss of mitogenic factors, serum starvation, drug treatment, etc.) (Stallaert et al. 2022). Thus, we find it very likely that additional G0 states with distinct transcriptional signatures will be identified. The ccAFv2 ANN and its associated training software are fully equipped to integrate these additional G0 states. Future studies that extend the cell cycle classifier to include novel G0 states holds immense potential for advancing our understanding of quiescence in biological systems. By leveraging advanced computational methods, high-throughput technologies, and interdisciplinary approaches, researchers can unravel the complexities of cellular dormancy and pave the way for innovative strategies to manipulate quiescent cell behavior to improve health and combat disease.

## Methods

### Culture of LGG glioma neurospheres

For the “no growth factors” condition, cells from LGG glioma neurospheres (LGG275, BT237) were dissociated, seeded into two poly-D-lysine and laminin coated T25 cm^2^ flasks at a density of 40,000 cells/cm^2^, and cultured for 4 days using medium without growth factors. For the “with growth factors” condition, cells were cultured for 4 days as neurospheres with EGF and FGF2 at 10µg/L and heparin at 2mg/L in PolyHEMA coated flasks, and medium was replaced 1 day before single cell sequencing.

### scRNA-seq library preparation and sequencing of LGG glioma neurospheres

Cells were dissociated and single cell suspensions loaded onto the Chromium controller (10x Genomics, Pleasanton, CA) to generate single-cell Gel Beads-in-Emulsion (GEMs). The single-cell RNA-seq libraries were prepared using the Chromium Next GEM Single Cell 3’ Reagent Kits V3.1 (Dual Index, P/N 1000268, 10x Genomics). Briefly, reverse transcription was performed at 53°C for 45 min followed by incubation at 85°C for 5 min. GEMs were then broken and the single-stranded cDNAs were cleaned up with DynaBeads MyOne Silane Beads (Thermo Fisher Scientific; P/N 37002D). The cDNAs were PCR amplified, cleaned up with SPRIselect beads (SPRI P/N B23318), fragmented, end-repaired, A-tailed, and size-selected with SPRIselect beads. Indexed adapters were ligated and cleaned up with SPRIselect beads. The resulting DNA fragments were PCR amplified and size selected with SPRIselect beads.

The size distribution of the resulting libraries was monitored using a Fragment Analyzer (Agilent Technologies, Santa Clara, CA, USA) and the libraries were quantified using the KAPA Library quantification kit (Roche, Basel, Switzerland). The libraries were denatured with NaOH, neutralized with Tris-HCl, and diluted to 150 pM. Clustering and sequencing were performed on a NovaSeq 6000 (Illumina, San Diego, CA, USA) using the paired-end 28-90 nt protocol on one lane of an SP flow cell and on one lane of an S4 flow cell. Sequencing data can be accessed from NCBI SRA. Both library preparation and sequencing were performed at the Montpellier GenomiX facility (MGX) in Montpellier, France.

### Data analysis

Image analyses and base calling were performed using the NovaSeq Control Software and the Real-Time Analysis component (Illumina). Demultiplexing was performed using the 10x Genomics software Cellranger mkfastq (v7.1.0), a wrapper of Illumina’s bcl2fastq (v2.20). The quality of the raw data was assessed using FastQC (v0.11.9) from the Babraham Institute and the Illumina software SAV (Sequencing Analysis Viewer). FastqScreen (v0.15.1) was used to identify potential contamination. Alignment, gene expression quantification and statistical analysis were performed using Cell Ranger count with the human’s transcriptome (GRCh38). To discard ambient RNA falsely identified as cells, Cell Ranger count was run a second time with the option--force-cells to force the number of cells to detect. Cell Ranger aggr was used to combine each sample results into one single analysis. Cell Ranger output files can be accessed from NCBI GEO at GSE263796.

### scRNA-seq, snRNA-seq, and ST-seq neuroepithelial datasets

In total 42 scRNA-seq, 11 snRNA-seq, and 8 ST-seq datasets were processed and employed in the studies used to characterize the ccAFv2 classifier. Detailed descriptions of the source, quality control, processing, normalization of each dataset can be found in the **Supplemental Material**. Analyses were conducted in R (R Core Team 2025) using the specified packages.

### Implementation of the ccAFv2 ANN model

The core algorithm of the ccAFv2 is a fully connected artificial neural network (ANN) implemented using the Keras API (v2.12.0) that employs TensorFlow (v2.12.0) to construct ANNs. A fully connected ANN model was developed to classify the cell cycle state of single cells (**Figure 1A**). The input for a single cell is expression for the 861 most highly variable genes (log_2_(FC) > 0.25; p value adj < 0.05) from O’Connor et al., 2021. A dense input layer takes in the expression of the 861 and is fully connected to the first hidden layer comprised of 600 neurons. The first hidden layer is fully connected to the second hidden layer comprised of 200 neurons which then connects to the output layer of seven neurons (one for each cell cycle class: Neural G0, G1, Late G1, S, S/G2, G2/M, and M/Early G1). A SoftMax regression function in the output layer is used to compute the likelihood for each class. Overfitting in the ANN is prevented through the incorporation of two dropout layers using a dropout rate of 50%. The first dropout layer is positioned between the first and second hidden layers and the second dropout layer is between the second hidden layer and the output layer (**Figure 1A**, Xie et al. 2019). Neuron activation functions were modeled using the Rectified Linear Unit (ReLU) function. The loss function for the ccAFv2 ANN was categorial cross-entropy and Stochastic Gradient Descent (SGD) was used to optimize the learning. The predicted class for a single cell is identified as the highest likelihood exceeding the specified threshold (**Figure 1B**). By default, the threshold is set at 0.5 and can be adjusted within the range of 0 to 1. If a cell’s likelihood falls below the threshold it is classified as “Unknown.”

### Training the ccAFv2 ANN classifier

The ccAFv2 ANN model was trained on the 2,692 cells and 861 genes from the U5-hNSCs dataset (O’Connor et al. 2021) using the labels from O’Connor et al., 2021. The training process encompassed ten epochs repeated five times consecutively. In each epoch, the training data was randomly partitioned into 80% for training and 20% for testing, with the testing subset held out to assess training accuracy.

### Comparing ccAFv2 to other classification methods

Classifiers were trained using the scRNA-seq gene expression of 2,962 cells with 861 genes and cell cycle labels from the U5-hNSCs. The ccAFv2 classifier was tested against: (i) support vector machine with reject option (SVMrej; classification cutoff ≥ 0.7), a general-purpose classifier from the Scikit-learn library; (ii) random forest (RF), another general-purpose classifier from the Scikit-learn library; (iii) *k*-nearest neighbor (KNN) from the SCANPY ingest method (Wolf et al., 2018); and (iv) neural network (NN) ACTINN (Ma & Pellegrini, 2020). Classifier performance was determined using F1 scores computed for each cell cycle state. Ten-fold cross-validation with an 80% training and 20% testing split was used to determine the variance of F1 scores for each cell cycle state from each classifier. A Student’s *t*-test was used determine if the mean of the F1 scores were significantly lower than ccAFv2.

### Optimizing the number of neurons in hidden layers

The configuration of neurons in the two hidden layers is designed to reduce the number of neurons at each layer from the 861 input genes down to the 7 cell cycle states. In total, 18 ccAFv2 models were trained using the U5-hNSCs dataset to determine the optimal number of neurons for these hidden layers. This involved testing at increments of 100 the number of neurons in the first hidden layer within the range of 200 to 700 neurons and in the second hidden layer within the range of 100 to 400 neurons. For comparisons the F1 scores were computed for each cell cycle state. Each model was also tested on pre-processed scRNA-seq data of glioma stem cells (BT322, BT324, BT326, BT333, BT363, BT368) and tumor cells (BT363, BT368) from Couturier et al. 2022 (**Supplemental Table S11**), along with Grade 2 Astrocytoma (LGG275; AUGUSTUS et al. 2021) (**Supplemental Table S12**). For these datasets, Adjusted Mutual Information (AMI) scores, with the reference labels derived from ccSeurat calls, and the number of cells predicted were calculated using the AMI function from the aricode package in R. Barcodes with an “Unknown” ccAFv2 label were removed before metrics were calculated.

### Computing feature importance

Feature importance for all 861 ccAFv2 features was determined by permuting each feature’s expression, running ccAFv2 with the permuted expression matrix, and comparing the likelihoods of all cells for a specific ccAFv2 state to the unpermuted likelihoods of the same cells. The average difference in likelihood was computed for each feature in each ccAFv2 state. A negative average difference in likelihood indicates that a feature was important, and the most negative features are the most important.

### Comparing ccAFv2 to existing cell cycle classifiers

The performance of ccAFv2 was compared with existing cell cycle state classifiers: ccAF (v1) (O’Connor et al. 2021), Seurat (Hao et al. 2021), Tricycle (Zheng et al. 2022), SchwabeCC (Schwabe et al. 2020; Zheng et al. 2022), reCAT (Liu et al. 2017), Peco (Hsiao et al. 2020) and Cyclone (Scialdone et al. 2015). Each classifier was applied to the PCW 9 R1 (Zeng et al, 2023) and BT322 (Couturier et al. 2020b) scRNA-seq datasets. Data was prepared as required to run each classifier method. The quality of predicted cell cycle states for each classification method was determined by computing the AMI score relative to reference cell cycle states. Ten-fold cross-validation with a 20% hold-out testing set was used to determine the variance of AMI scores for each cell cycle state from each classification method. For both datasets, the ccSeurat predicted cell cycle states were used as the reference for computing AMI scores. Cells with “Unknown” labels were excluded when computing AMI scores. The median AMI scores were tabulated and plotted against the number of predicted states for each classifier. Representative cell cycle state predictions for each classification method were also visualized as UMAPs.

Because each classifier predicts different numbers of cell cycle states (3 – 8 cell cycle states) it was necessary to use simulated datasets to determine the range of AMI scores that correspond to specific amounts of similarity to the reference. Predicted cell cycle states with 3 to 8 states were simulated that contained specific 0 to 100% similarity to a simulated reference, at 10% increments. The average AMI from 100 simulated cell cycle classifications was computed for each specific amount of similarity to a simulated reference and plotted as a guide to assess the quality between classification methods with different numbers of cell cycle states.

### Finding the optimal likelihood threshold

A neuroepithelial dataset of *in vivo* hNSCs from fetal tissue at 3 to 12 weeks post-conception from Zeng et al., 2023, that was independent of the ccAFv2 training data, was used to determine the optimal likelihood threshold. Random sub-sampling of 90% of cells for each timepoint was used to determine the variance of the classifications and ccAFv2 was applied with likelihood thresholds ranging from 0.0 to 0.9 by increments of 0.1. For each iteration metrics were collected including the number of cells predicted, and an AMI score computed using ccSeurat cell cycle states as the reference. Cells with “Unknown” labels were excluded when computing AMI scores. Metrics were not computed when 20 or fewer cells were predicted.

Student’s *t*-tests were used to compare AMIs computed at each examined likelihood threshold with those derived from a likelihood threshold of 0.0, which is equivalent to not using a likelihood threshold, and a significant difference was considered a p-value ≤ 0.05. A baseline for comparison was provided by random removal of an equivalent percentage of cells that were classified as “Unknown” for each likelihood threshold, and an AMI was computed with the remaining cells. Student’s *t*-tests were used to compare AMIs of the likelihood thresholded and random removal at each likelihood threshold, and a significant difference was considered a p-value ≤ 0.05.

### Cell cycle state validation using hNSCs (PCW 9 R1)

We used the hNSCs collected from whole fetal brain at nine weeks post-conception replicate 1 (PCW 9 R1) to validate the cell cycle states assigned by ccAFv2. After quality control (**Supplemental Table S13**) and normalization with sctransform, 5575 cells were classified into distinct cell cycle states using ccAFv2. We selected five key markers of cell cycle states: *CCND1* (Late G1), *CCNE2* (S), *CCNA2* (G2/M), *CCNB1* (G2/M), and *CDK1* (G2/M) to assess the expression patterns associated with these phases. The average expression levels of the genes were calculated and visualized using violin plots, which were grouped according to the cell cycle states predicted by ccAFv2. In addition, we monitored the dynamic changes in the average expression of each key marker as cells transitioned between different cell cycle states. Student’s *t*-tests were used to determine if the marker expression was significantly different at each cell cycle state compared to the G1 state. Finally, relative expression levels of top marker genes for each cell cycle state were identified using FindAllMarkers() and visualized using a heatmap, with cells grouped by cell cycle state.

### Comparison of Neural G0 state with G0 arrest signature using QuieScore

We applied the QuieScore algorithm (https://github.com/dkornai/QuieScore) to the U5-hNSCs using the cancer type parameter of “LGG”. The G0 cells were identified by a q_score_raw of greater than 3. We evaluated the similarity between the QuieScore-identified G0 cells with the ccAFv2-identified Neural G0 cells using hypergeometric enrichment analysis.

### Determining the sensitivity of ccAFv2 to missing genes

Sensitivity analysis was conducted on the U5-hNSC dataset by randomly setting a defined percentage of classifier genes (1-90%) to zero and applying the ccAFv2 classifier. Each percentage of classifier genes was subsampled ten times and for each iteration the metrics error rate and percentage of cells predicted were recorded.

### Demonstrating the generalizability of ccAFv2

The 245,906 human fetal cells 3 to 12 weeks post conception (Zeng et al., 2023) encompassing fifteen cell types that represent all three germ layers (**Supplemental Table S14**) were classified by ccAFv2. Positive marker genes for the Neural G0 cells were identified for each cell type using the FindAllMarkers function (log_2_ fold change ≥ 0.25; adjusted p-value ≤ 0.05). The Neural G0 markers were tabulated among each dataset and across all datasets to identify common Neural G0 marker genes.

### Regressing out cell cycle transcriptional signatures using ccAFv2 marker genes

The average expression from the marker genes for each cell cycle state (**Supplemental Table S9**) was computed using the AddModuleScore function in Seurat. The S and G2/M or Late G1, S, S/G2, G2/M, M/Early G1 module scores were regressed out in the SCTransform function in Seurat. The variance explained by the first principal component of the marker genes was used as a metric for co-expression of the cell cycle transcriptional signatures. Empirical p-values were calculated by comparing the observed variance explained to the variance explained of 1,000 randomly sampled gene sets of the same size. Significantly regressing out the cell cycle transcriptional signature was determined by a reduction in the variance explained that made the empirical p-value non-significant (>0.05).

### Application of ccAFv2 to neuroepithelial scRNA**-**seq and snRNA-seq profiling studies

To maximize overlap with the ccAFv2 input genes, we enabled the option to apply SCTransform (do_sctransform) for SCTransformed datasets. The species (‘human’ or ‘mouse’) and gene ID (‘Ensembl’ or ‘symbol’) options were configured based on the specifications of each dataset.

Predicted cell cycle states were collected from each dataset and integrated with meta information.

### Application of ccAFv2 to ST-seq data

We downloaded the transcriptome profiles for a 5 μm section of a male C57BL/6 mouse embryo taken from an FFPE tissue block obtained from Charles River Laboratories that was made public by 10x Genomics (https://www.10xgenomics.com/datasets/visium-hd-cytassist-gene-expression-libraries-of-mouse-embryo). The 10x Visium HD Gene Expression Library preparation kit afforded a resolution of 2 μm^2^ spots and details about sample preparation and library performance and QC can be found on the 10x website linked above. In Seurat the 2 μm^2^ spots were binned into 8 μm^2^ bins, the data log normalized, and ccAFv2 was applied to predict cell cycle states for each spot. Expression of key genes was plotted using the normalized and scaled values.

### R and Python package for ccAFv2

The ccAFv2 classifier has been implemented as an R package (https://github.com/plaisier-lab/ccafv2_R) that can be installed and used as part of a Seurat workflow, and works for both Seurat version 4 and 5 (**Supplemental Figure S14**). Due to differences in the Seurat v5 SCTransform function it was necessary to set the vst.flavor equal to “v1” to make it equivalent to Seurat v4.3.0.1, and leaving the vst.flavor as the default in v5 leads to only small differences (**Supplemental Figure S14**). For the Seurat v5.0.2 the matrixStats package was required to be v1.1.0. Additionally, the ccAFv2 classifier has been implemented as a Python PyPI installed package (https://pypi.org/project/ccAF/) that can be installed and used as part of a SCANPY workflow. It should be noted that SCTransform normalization is the suggested method for preparing data that will be classified by ccAFv2, and as of now there is no SCTransform option in SCANPY.

### Culture of human skeletal muscle satellite cells (hSkMSCs)

hSkMSCs were purchased from ScienCell Research Laboratories (P/N 3510, ScienCell) and were grown in Skeletal Muscle Cell Medium (P/N 3501; ScienCell) on Nunclon Delta-treated cell culture flasks and passaged according to vendor protocols. Cells were detached from their plates using Trypsin/EDTA Solution (P/N 183; ScienCell) and collected with Trypsin Neutralizing Solution (P/N 113; ScienCell).

### scRNA-seq characterization of hSkMSCs

hSkMSCs were grown up to 80% confluency, washed with Molecular Biology Grade PBS (P/N 45001-130, VWR), dissociated with Trypsin/EDTA Solution, and collected in Trypsin Neutralizing Solution. After centrifugation at 300 x g for 5 minutes, cells were resuspended in Molecular Biology Grade PBS containing 0.04% BSA and counted using an automated cell counter. Cells were then diluted to 1,000 cells/μl. scRNA-seq library preparation was performed by the ASU Genomics Core facility. Samples were processed using the 10x Chromium Single Cell 3′ Gene Expression v3.1 kit into a single library (10x Genomics). The quality of the library was determined using Agilent TapeStation automated electrophoresis. Samples were sequenced at an average read depth of 100,000 reads per cell (Illumina, Novogene). The 10x Genomics CellRanger v7.0.1 was used to align to the Human reference genome GRCh38-2020- A (GRCh38), quantify, and provide basic quality control metrics for the scRNA-seq data. The 10x CellRanger outputs for 7,795 hSkMSCs was loaded into Seurat. Filtering and downstream analyses was done using quality control and downstream processing code templates provided in https://github.com/plaisier-lab/ccAFv2. Standard Seurat filters were applied requiring that the cells had to have a least 200 features per cell, and transcripts need to be expressed in at least 3 cells. Then the cells were further filtered to 7,207 hSkMSCs by requiring the number of UMIs per cell to fall within the range of 4,000 to 100,000, and the percentage of mitochondrial genes expressed relative to total expression per cell was required to fall within the range of 0.9 to 10%. The filtered cells were then normalized using SCTransform (Hafemeister and Satija 2019), principal components were calculated, and a UMAP was generated.

### Staining for quiescent-like G0 cells

This staining protocol is based on a protocol developed by Gookin et al., 2017 (Gookin et al. 2017) to identify a G0/quiescent subpopulation and has been adapted for fluorescence-activated cell sorting (FACS) and to preserve RNA integrity. Cells were expanded in Nunclon Delta-treated cell culture flasks to achieve the desired cell count, accounting for a 50% loss during staining preparation before downstream FACS.

<u>Replication stain</u>: Replicating cells were labeled with a synthetic nucleotide Tetramethylrhodamine-dUTP (P/N 17023, AAT Bioquest) transported into the cells using a synthetic nucleotide triphosphate transporter (SNTT) the BioTracker NTP-Transporter Molecule (P/N SCT064, Millipore Sigma) (Zawada et al. 2018; Gookin et al. 2017). When cells reached 70% confluence, they were washed with tricine buffer, the SNTT and synthetic nucleotide were diluted so each component was 20 µM in tricine buffer, added to cells, and incubated at 37°C and 5% CO2 for 5 minutes to transport fluorescently labeled synthetic nucleotide into the cells. The stain was then aspirated and replaced with complete culture medium, and the cells were incubated at 37°C and 5% CO2 for 1 hour to allow time for replicating cells to incorporate the fluorescently labeled synthetic nucleotides into their genome’s.

<u>Viability stain</u>: Cells were then washed with Molecular Biology Grade PBS, dissociated with Trypsin/EDTA Solution, and collected in Trypsin Neutralizing Solution. After centrifugation at 300 x g for 5 minutes, cells were resuspended in Molecular Biology Grade PBS and counted using an automated cell counter. Cells were centrifuged again, PBS was removed, resuspended in 1:1000 LIVE/DEAD™ Fixable Near-IR Dead Cell Stain Kit (P/N L34975, ThermoFisher Scientific) using manufacturer instructions, and incubated for 30 minutes at room temperature in the dark.

<u>Fixation and rehydration</u>: Cells were washed with 0.5% Ultra-Pure BSA (P/N AM2616, ThermoFisher Scientific) in Molecular Biology Grade PBS and centrifuged at 300 x g two times. The cells were fixed by first resuspending them in ice-cold Molecular Biology Grade PBS at a volume of 200 µl per 1 million cells. Then, ice-cold 100% methanol was added dropwise at a volume of 800 µl per 1 million cells, with gentle shaking. Cells were then incubated for at least 30 minutes at-20°C. After fixation, cells were kept on ice. Cells were rehydrated with cold 3X SSC Rehydration Cocktail (Chen et al. 2018), followed by centrifugation at 500 x g for 5 minutes. Cells were washed one more time with the SSC Rehydration Cocktail, and one time with 0.5% Ultra-Pure BSA in Molecular Biology Grade PBS.

<u>Phosphorylated RB (pRB) staining</u>: Hypo-phosphorylation of pRB is an established indicator of a cell being in a quiescent G0 state (Gookin et al. 2017). Primary antibody for pRB (Ser807/811) (P/N 8516T, Cell Signaling Technologies) was added at a dilution of 1:200 in 0.5% Ultra-Pure BSA in Molecular Biology Grade PBS and incubated overnight at 4°C. Cells were washed three times with 0.5% Ultra-Pure BSA in Molecular Biology Grade PBS and then fluorescently labeled secondary antibody (P/N 4412, Cell Signaling) was added at a dilution of 1:1000 dilution for 30 minutes. Samples were then washed two times with 0.5% Ultra-Pure BSA in Molecular Biology Grade PBS.

<u>DNA staining</u>: The ploidy of cells was determined using Hoechst DNA stain (Gookin et al. 2017). Prior to FACS cells were stained with 2 ug/ml of Hoechst DNA stain (P/N 561908, BD) diluted in Molecular Biology Grade PBS, without BSA.

### Fluorescence-activated cell sorting of G0 cells

Cells were filtered using sterile CellTrics 30 μm filters (P/N 04-004-2326, Sysmex) into sterile, nuclease-free 5 ml polystyrene round-bottom tubes for sorting (P/N 352235, Corning) and kept covered from light and on ice until sorting. Cells were stained using the following experimental design to define gates and have adequate controls: 1) Hoechst only, 2) Live/Dead only, 3) replication only, 4) pRB only, 5) replication fluorescence minus one (FMO), 6) pRB FMO, and 7) all stains. Example of gating can be seen in **Figure 4A**. Cell sorting was performed using the FACSymphony flow cytometer (BD). G0 cells were defined as viable cells, that were diploid (2N), with low pRB. FACS data analysis was performed using FlowJo (BD).

### RNA-sequencing of G0 cells

RNA was extracted from 400,000 sorted cells from two biological replicates using the Qiagen RNeasy Micro Kit (P/N 74004, Qiagen). The concentration and quality of RNA was determined by Nanodrop (Thermo Scientific) and High Sensitivity RNA TapeStation (Agilent). Both G0 samples had more than 300 ng of RNA and RIN scores of greater than 9. Samples were sent for sequencing on the NovaSeq X Plus (Illumina, Novogene). A docker RNA-seq pipeline (cplaisier/star_2_7_1a_grch38_p21; DOI = https://doi-org.ezproxy1.lib.asu.edu/10.5281/zenodo.5519663) was employed to align reads from FASTQ files to the genome using STAR v2.7.1a (Dobin et al. 2013) and GENCODE genome build GRCh38 Release 31 (Frankish et al. 2023). Counts were tabulated using htseq-count (Putri et al. 2022). DESeq2 (Love, Huber, and Anders 2014) was used for subsequent differential gene expression analysis.

### Correlating RNA-seq and scRNA-seq data

DESeq2-normalized RNA-seq data and sctransform-normalized scRNA-seq data were loaded into R. Marker genes were selected by identifying highly variable scRNA-seq genes with more than 10 counts in the bulk G0 subpopulations. Additionally, 861 ccAFv2 marker genes present in the RNA-seq data were included. Both scRNA-seq and RNA-seq datasets were filtered to include these 3,120 marker genes. The expression profiles of individual cells in the scRNA-seq data were correlated with the G0 RNA-seq profiles using the corr package in R (Makowski et al., 2020), using the Spearman method.

## Data Access

All raw and processed sequencing data for the hSkMSC FACs sorted RNA-seq and scRNA-seq generated in this study have been submitted to NCIB Gene Expression Omnibus (GEO; https://www.ncbi.nlm.nih.gov/geo/) under accession number GSE285220. All raw and processed sequencing data for the four LGG scRNA-seq generated in this study have been submitted to GEO under accession number GSE263796.

All other data used in our analyses are available on Zenodo (https://zenodo.org/doi/10.5281/zenodo.10963136). We also provide all code on github.com (https://github.com/plaisier-lab/ccAFv2) and Docker images on DockerHub that were used to run all analyses (https://hub.docker.com/r/cplaisier/ccafv2_extra and https://hub.docker.com/r/cplaisier/ccnn).

### R package for ccAFv2

We have developed an R package that can be installed using devtools from github. The instructions for installation and usage can be found on github: https://github.com/plaisier-lab/ccafv2_R

### Python package for ccAFv2

We have also developed a Python package that can be installed using pip from PyPi. The instructions for installation and usage can be found on PyPi and github: https://pypi.org/project/ccAFv2/ and https://github.com/plaisier-lab/ccAFv2_py

### Docker images for ccAFv2

We also provide Docker images that include all dependencies and ccAFv2 preinstalled to make the package more user friendly. Please see the github repositories for information about how to get, run, and use the Docker images.

‒ R package:
`o` Seurat v4: https://hub.docker.com/r/cplaisier/ccafv2_seurat4
`o` Seurat v5: https://hub.docker.com/r/cplaisier/ccafv2_seurat5
‒ Python package: https://hub.docker.com/r/cplaisier/ccafv2_py

## Acknowledgments

This work was supported by the following grants: NINDS/NIH (R01NS119650: A.P., P.P., C.L.P.) and (R01NS123038: CLP); NCI/NIH (R01CA190957; R21CA170722; P30CA15704: P.P.); DoD Translational New Investigator Award (CA100735: P.P.); and the Pew Biomedical Scholars Program (P.P.). The authors acknowledge Joy Blain, Anna Engelbrektson, and Adam Kindelin for their assistance in scRNA-seq library preparation, quality control, and FACS. The BT237 cell line was provided by Keith Ligon from the Dana-Farber Cancer Institute Center for Patient Derived Models. Some components of Figure 5 were made using Biorender.com.

## Author contributions

Project conception and experimental design were carried out by CLP, SAO, PJP, and AP. Implementation of ccAFv2 and the R and Python packages was performed jointly by SAO, RH and CLP. scRNA-seq of LGG primary cell lines was performed by LG and J-PH. Single cell/nuclei dataset curation was performed by SAO under the supervision of CLP. Testing of ccAFv2 was performed by SAO under the supervision of CLP. ST-seq dataset curation and testing was performed by CLP. Interpretation of ST-seq results was performed by BBB and CLP. The manuscript was written by SAO, BBB, and CLP with input from all authors.

