## Supplementary Materials for "Classifying cell cycle states and a quiescent-like G0 state using single-cell transcriptomics"

### Table of Contents


**Supplemental Methods** **3**

*Human neural stem cells* 3

*Glioblastoma tumor and cancer stem cells* 3

*Low grade glioma IDH mutant tumor cells* 3

*Human neural stem cells from fetal tissue at post-conception weeks 3-12* 3

*Human cells from fetal tissue at post-conception weeks 3-12* 4

*Atlas of the developing human telencephalon* 4

*Atlas of adult neurogenesis in the ventricular-subventricular zone (V-SVZ)* 4

*Activated and quiescent neural stem cells* 4

*Atlas of developing human spinal cord* 5

**Supplemental Tables** **6**

*Supplemental Table S1* 6

*Supplemental Table S2* 6

*Supplemental Table S3* 6

*Supplemental Table S4* 7

*Supplemental Table S5* 7

*Supplemental Table S6* 7

*Supplemental Table S7* 7

*Supplemental Table S8* 7

*Supplemental Table S9* 8

*Supplemental Table S10* 8

*Supplemental Table S11* 8

*Supplemental Table S12* 8

*Supplemental Table S13* 9

*Supplemental Table S14* 10

*Supplemental Table S15* 10

**Supplemental Files** **11**

**Supplemental Figures** **12**

*Supplemental Figure S1* 12

*Supplemental Figure S2* 14

*Supplemental Figure S3* 16

*Supplemental Figure S4* 17

*Supplemental Figure S5* 19

***Supplemental Figure S6*** *20*
***Supplemental Figure S7*** *21*
***Supplemental Figure S8*** *22*
***Supplemental Figure S9*** *23*
***Supplemental Figure S10*** *24*
***Supplemental Figure S11*** *25*
***Supplemental Figure S12*** *26*
***Supplemental Figure S13*** *27*
***Supplemental Figure S14*** *28*

### Supplemental Methods

#### *Neuroepithelial datasets*

**Human neural stem cells**

The 10X Cell Ranger output for 3,049 U5-hNSCs (O'Connor et al. 2021) (available on NCBI GEO as GSM3267241 from GSE117003) was loaded into Seurat. Standard Seurat filters were applied requiring that the cells had to have at least 200 features per cell, and transcripts need to be expressed in at least three cells. Then the cells were further filtered to 3,035 cells by requiring the number of UMIs per cell to fall within the range of 1,000 to 20,000 UMIs, and the mitochondrial percentage of genes expressed per cell to fall within the range of 0.01 to 7%. For the purposes of these studies the 73 G1/other cells were removed because they did not align with any known cell cycle state. The 2,962 U5-hNSC cells were then normalized using SCTransform (Hafemeister and Satija 2019), principal components were calculated, and a UMAP was generated. Labels were transferred from O'Connor et al., 2021.

**Glioblastoma tumor and cancer stem cells**

The 10X Cell Ranger outputs for two tumors (BT363 and BT368) and five cancer stem cell lines (BT322, BT324, BT333, BT363, and BT368) derived from tumors (available on the European Genome-Phenome Archive as EGAS00001004422) were loaded into Seurat. Standard Seurat filters were applied requiring that the cells had to have at least 200 features per cell, and transcripts need to be expressed in at least three cells. Each sample was further filtered by requiring the number of UMIs per cell to fall within a range of UMIs (**Supplemental Table S11**), and the mitochondrial percentage of genes expressed per cell to fall within a range of percents (**Supplemental Table S11**). The cells from each sample were then normalized using SCTransform (Hafemeister and Satija 2019), principal components were calculated, and a UMAP was generated. Clustering and expression of marker genes was used to remove non-tumor cell types from each sample:  oligodendrocytes - *MBP* and *PLP1* (Valério-Gomes et al. 2018); astrocytes - *ETNPPL* (Zhang et al. 2016); neurons - *RBFOX* (Herculano-Houzel and Lent 2005); or immune cells - *AIF1*, *CD14*, *CX3CR1*, *PTPRC*. The number of cells for each sample are listed in the tumor cells column of Supplemental Table S11. The ccAFv2 classifier was applied to determine the cell cycle state of each cell.

**Low grade glioma IDH mutant tumor cells**

The 10X Cell Ranger outputs for LGG275 (a grade 2 astrocytoma) (Augustus et al. 2021) and BT237 (a grade 3 oligodendroglioma) (now available on NCBI GEO as GSE263796 were loaded into Seurat. Standard Seurat filters were applied requiring that the cells had to have at least 200 features per cell, and transcripts need to be expressed in at least three cells. Each sample was further filtered by requiring the number of UMIs per cell to fall within a range of UMIs (**Supplemental Table S12**), and the mitochondrial percentage of genes expressed per cell to fall within a range of percents (**Supplemental Table S12**). The remaining cells were then normalized using SCTransform (Hafemeister and Satija 2019), principal components were calculated, and a UMAP was generated. The ccAFv2 classifier was applied to determine the cell cycle state of each cell.

**Human neural stem cells from fetal tissue at post-conception weeks 3-12**

The data for the human gastrulation and early brain development scRNA-seq atlas from Zeng et al., 2023 (Zeng et al. 2023) (available from NCBI GEO as GSE155121) was downloaded as an h5ad file. The cells were subset to only the hNSCs by filtering the 'Main Cell Type', which is a metadata column in the h5ad object, for the keyword 'NSC' using the Python package scanpy

(Wolf et al. 2018). Next, the 109,578 hNSCs were loaded into Seurat. Standard Seurat filters were applied requiring that the cells had to have at least 200 features per cell, and transcripts need to be expressed in at least three cells. Then the cells were subset by ‘week stage’ and filtered by requiring the number of UMIs per cell to fall within a range of UMIs (**Supplemental Table S13**), and the mitochondrial percentage of genes expressed per cell to fall within a range of percents (**Supplemental Table S13**). The remaining cells were then normalized using SCTransform (Hafemeister and Satija 2019) and the ccAFv2 classifier was applied to determine the cell cycle state of each cell.

**Human cells from fetal tissue at post-conception weeks 3-12**

The data for the human gastrulation and early brain development scRNA-seq atlas from Zeng et al., 2023 (Zeng et al. 2023) (available from NCBI GEO as GSE155121) was downloaded as an h5ad file. The cells were subset to each ‘Main Cell Type’, which is a metadata column in the h5ad object, using the Python package scanpy (Wolf et al. 2018) and saved as individual h5ad files. Data were loaded into Seurat. Standard Seurat filters were applied requiring that the cells had to have at least 200 features per cell, and transcripts need to be expressed in at least three cells. Then the cells were subset by ‘week stage’ and filtered by requiring the number of UMIs per cell to fall within a range of UMIs, and the mitochondrial percentage of genes expressed per cell to fall within a range of percents (**Supplemental Table S14**). The remaining cells were then normalized using SCTransform (Hafemeister and Satija 2019) and the ccAFv2 classifier was applied to determine the cell cycle state of each cell. Positive marker genes (log2 fold change ≥ 0.25; adjujsted p-value ≤ 0.05) were identified for each cell cycle state using the FindAllMarkers function. Neural G0 markers were tabulated among each dataset and across all datasets to identify common Neural G0 marker genes.

**Atlas of the developing human telencephalon**

The processed data for the 4,261 cells from the Nowakowski et al., 2017 (Nowakowski et al. 2017) scRNA-seq atlas of the developing human telencephalon was downloaded from http://bit.ly/cortexSingleCell and loaded into Seurat. The ccAFv2 classifier was applied to determine the cell cycle state of each cell, without running SCTransform.

**Atlas of adult neurogenesis in the ventricular-subventricular zone (V-SVZ)**

The processed and SCTransformed scRNA-seq data for 24,261 cells from the ventricular-subventricular zone (V-SVZ) of adult mice from Cebrian-Silla et al., 2021 (Cebrian-Silla et al. 2021) was loaded into Seurat as an RDS file (available from GSM5039270 from NCBI GEO GSE165554). The ccAFv2 classifier was applied to determine the cell cycle state of each cell.

**Activated and quiescent neural stem cells**

The processed data for the 129 cells from the Llorens-Bobadilla et al., 2015 (Llorens-Bobadilla et al. 2015) scRNA-seq characterization of activated and quiescent NSCs was loaded into Seurat as an expression matrix (available from NCBI GEO GSE67833). In lieu of SCTransform, the data was log-normalized, 3,000 variable features were detected, and the data was then scaled. The ccAFv2 classifier was applied to determine the cell cycle state of each cell, without running SCTransform.

The raw count matrix for the 279 cells from the Dulken et al., 2017 (Dulken et al. 2017) (available as NCBI BioProject PRJNA324289) were loaded into Seurat. Standard Seurat filters were applied requiring that the cells had to have at least 200 features per cell, and transcripts need to be expressed in at least three cells. The remaining cells were then normalized using

SCTransform (Hafemeister and Satija 2019). The ccAFv2 classifier was applied to determine the cell cycle state of each cell.

**Atlas of developing human spinal cord**

The 10X Cell Ranger outputs for eleven samples with both scRNA-seq and snRNA-seq from an atlas of the developing human spinal cord by Zhang et al., 2021 (Zhang et al. 2021) across 8 to 23 weeks post-conception (8 – 23 PCW; available on NCBI GEO as GSE136719) were loaded into Seurat. Standard Seurat filters were applied requiring that the cells/nuclei had to have at least 200 features per cell/nuclei, and transcripts need to be expressed in at least three cells/nuclei. Each sample was further filtered by requiring the number of UMIs per cell/nuclei to fall within a range of UMIs in the table below (**Supplemental Table S15**), and the mitochondrial percentage of genes expressed per cell to fall within a range of percents or for nuclei the counts per nuclei within a range of counts (**Supplemental Table S15**). The cells/nuclei from each sample were then normalized using SCTransform (Hafemeister and Satija 2019), principal components were calculated, and a UMAP was generated. The ccAFv2 classifier was applied to determine the cell cycle state of each cell.

### Supplemental Tables

**Supplemental Table S1**. ccSeurat calls versus Hoechst and FUCCI-defined ground truth labels.

| | ccSeurat | | |
|---|---|---|---|
| **Hoechst** | **G1** | **S** | **G2M** |
| **G1** | 25 | 65 | 6 |
| **S** | 12 | 53 | 31 |
| **G2M** | 1 | 9 | 86 |

| | ccSeurat | | |
|---|---|---|---|
| **FUCCI** | **G1** | **S** | **G2M** |
| **G1** | 0 | 41 | 50 |
| **S** | 18 | 59 | 3 |
| **G2** | 2 | 10 | 64 |

**Supplemental Table S2**. Marker genes for each cell cycle state for U5-hNSCs.

[Supplemental Table S2 provided as a separate file]

**Supplemental Table S3**. Optimizing number of neurons in hidden layers.

[Supplemental Table S3 provided as a separate file]

**Supplemental Table S4**. Existing cell cycle classifiers.

| Package | Phases predicted | Model | Underlying data | Reference |
|---|---|---|---|---|
| ccAF | Neural G0, G1, Late G1, S, S/G2, G2/M, M/Early G1 | Neural network | Human neural stem cells | **Our work**: O'Connor et al, 2021 |
| Seurat/Scanpy | G1, S, G2/M | Gene scoring | Mouse hematopoietic progenitors | Butler et al, 2018 |
| tricycle | G1/G0, S, G2/M, M | Transfer learning | Mouse cortical neurospheres | Zheng et al, 2022 |
| SchwabeCC / Revelio | G1.S, S, G2, G2.M, M.G1 | PCA | mNeurosphere | Schwabe et al, 2020 |
| reCAT | G1, G1S, S, G2, G2M, M | Consensus traveling salesman problem and hidden Markov | Mouse embryonic stem cells | Liu et al, 2017 |
| peco | G1/G0, S, G2/M, M | Partition around medoids | Induced pluripotent stem cells | Hsiao et al, 2020 |
| cyclone | G1, S, G2M | Pairs method | Mouse embryonic stem cells | Scialdone et al, 2015 |

**Supplemental Table S5**. Likelihood threshold analysis for GSE155121.

[Supplemental Table S5 provided as a separate file]

**Supplemental Table S6**. Missing genes analysis for U5-hNSCs and PCW8 hNSCs.

[Supplemental Table S6 provided as a separate file]

**Supplemental Table S7**. Missing genes analysis for U5-hNSCs broken down by cell cycle state.

[Supplemental Table S7 provided as a separate file]

**Supplemental Table S8**. hSkMSC G0 marker genes.

[Supplemental Table S8 provided as a separate file]

**Supplemental Table S9**. ccAFv2 regression analysis.

[Supplemental Table S9 provided as a separate file]

**Supplemental Table S10**. Nowakowski et al. 2017 ccAF and ccAFv2 predictions.

[Supplemental Table S10 provided as a separate file]

**Supplemental Table S11.** Glioblastoma tumor and cancer stem cells QC filters.

| Sample | UMI bot. | UMI top | Mito. % bot. | Mito. % top | Resulting number of cells | Tumor cells |
|---|---|---|---|---|---|---|
| BT363 (Tumor) | 5,000 | 40,000 | 0.1 | 8 | 5,128 | 4,365 |
| BT368 (Tumor) | 6,000 | 50,000 | 0.1 | 8 | 1,556 | 1,262 |
| BT322 | 4,000 | 62,000 | 0.9 | 10 | 3,409 | 3,409 |
| BT324 | 5,000 | 40,000 | 0.9 | 6 | 4,377 | 3,351 |
| BT333 | 10,000 | 90,000 | 0.9 | 8 | 2,457 | 2,457 |
| BT363 | 5,000 | 80,000 | 0.9 | 10 | 4,917 | 4,770 |
| BT368 | 3,000 | 35,000 | 0.9 | 8 | 4,535 | 3,672 |

**Supplemental Table S12.** Low grade glioma IDH mutant tumor cells QC filters.

| Sample | UMI bot. | UMI top | Mito. % bot. | Mito. % top | Resulting number of cells |
|---|---|---|---|---|---|
| LGG275_GF | 5,000 | 76,000 | 0.1 | 15 | 2,232 |
| LGG275_noGF | 6,000 | 35,000 | 0.1 | 8 | 1,744 |
| BT237_GF | 6,000 | 90,000 | 1 | 15 | 1,485 |
| BT237_noGF | 6,000 | 90,000 | 3 | 17 | 3,074 |

**Supplemental Table S13.** Human neural stem cells from fetal tissues at post-conception weeks 3-12 QC filters.

| Sample | UMI bot. | UMI top | Mito. % bot. | Mito. % top | Resulting number of cells |
|---|---|---|---|---|---|
| PCW3-1 | 2,000 | 40,000 | 0.1 | 4 | 1,177 |
| PCW4-1 | 4,000 | 40,000 | 0.1 | 8 | 5,297 |
| PCW4-2 | 4,000 | 32,000 | 0.1 | 6 | 12,458 |
| PCW4-3 | 4,000 | 28,000 | 0.1 | 6 | 12,269 |
| PCW5-1 | 4,000 | 32,000 | 0.1 | 5 | 6,126 |
| PCW5-2 | 4,000 | 30,000 | 0.5 | 14 | 10,009 |
| PCW5-3 | 4,000 | 35,000 | 0.5 | 6 | 9,133 |
| PCW6-1 | 4,000 | 32,000 | 0.1 | 4 | 6,040 |
| PCW7-1 | 4,000 | 30,000 | 0.1 | 5 | 15,246 |
| PCW8-1 | 4,000 | 23,000 | 0.1 | 4 | 2,562 |
| PCW9-1 | 4,000 | 23,000 | 0.1 | 4 | 5,575 |
| PCW9-2 | 4,000 | 25,000 | 0.1 | 5 | 4,277 |
| PCW12-1 | 4,000 | 20,000 | 0.1 | 4 | 4,128 |

**Supplemental Table S14**. Cells (not including NSCs) from fetal tissues at post-conception weeks 3-12 QC filters.

[Supplemental Table S14 provided as a separate file]

**Supplemental Table S15.** Atlas of developing human spinal cord QC filters.

| Sample | Location | Cells or Nuclei | Raw cells | UMI bot. | UMI top | Mito. % or counts bot. | Mito. % or count top | Resulting number of cells |
|---|---|---|---|---|---|---|---|---|
| GW8 | Spinal Whole | Cells | 24,874 | 3,000 | 50,000 | 0.1 | 8 | 11,047 |
| GW8 | Spinal Whole | Nuclei | 25,282 | 200 | 15,000 | 200 | 6,000 | 25,269 |
| GW10 | Cervical | Cells | 13,907 | 3,000 | 50,000 | 0.1 | 8 | 5,763 |
| GW10 | Cervical | Nuclei | 7,509 | 200 | 55,000 | 200 | 10,000 | 7,485 |
| GW10 | Lumbar | Cells | 10,456 | 3,000 | 45,000 | 0.1 | 8 | 4,695 |
| GW10 | Lumbar | Nuclei | 19,887 | 200 | 70,000 | 200 | 11,000 | 19,869 |
| GW10 | Thoracic | Cells | 11,229 | 3,000 | 40,000 | 0.1 | 8 | 5,307 |
| GW10 | Thoracic | Nuclei | 7,026 | 200 | 55,000 | 200 | 10,000 | 7,011 |
| GW11 | Cervical | Cells | 18,691 | 3,000 | 50,000 | 0.1 | 5 | 11,377 |
| GW11 | Cervical | Nuclei | 9,069 | 200 | 15,000 | 200 | 5,500 | 9,067 |
| GW11 | Lumbar | Cells | 8,099 | 2,000 | 18,000 | 0 | 5 | 500 |
| GW11 | Lumbar | Nuclei | 8,099 | 200 | 15,000 | 200 | 6,000 | 8,094 |
| GW11 | Thoracic | Cells | 8,026 | 2,000 | 12,000 | 0 | 10 | 1,122 |
| GW11 | Thoracic | Nuclei | 17,763 | 200 | 30,000 | 200 | 6,000 | 17,676 |
| GW20 | Cervical | Cells | 11,182 | 3,000 | 30,000 | 0.1 | 8 | 9,441 |
| GW20 | Cervical | Nuclei | 10,146 | 200 | 18,000 | 200 | 6,000 | 10,137 |
| GW20 | Lumbar | Cells | 12,189 | 3,000 | 40,000 | 0.1 | 10 | 9,011 |
| GW20 | Lumbar | Nuclei | 8,493 | 200 | 16,000 | 200 | 6,000 | 8,484 |
| GW20 | Thoracic | Cells | 11,745 | 3,000 | 25,000 | 0.8 | 8 | 9,694 |
| GW20 | Thoracic | Nuclei | 10,004 | 200 | 12,000 | 200 | 5,000 | 9,999 |
| GW23 | Spinal Whole | Cells | 11,293 | 3,000 | 28,000 | 0.8 | 10 | 8,602 |
| GW23 | Spinal Whole | Nuclei | 11,293 | 200 | 28,000 | 200 | 5,000 | 11,217 |

### Supplemental Files

1. Supplementary_Code.tgz – This file contains the code for running data and analysis pipelines for producing paper figures. It can also be found at https://github.com/plaisier-lab/ccAFv2/tree/main/code. The data can be found at https://zenodo.org/doi/10.5281/zenodo.10963136.

### Supplemental Figures

**Supplemental Figure S1**. Building ccAFv2 and comparison of cell labels and ccAFv2 cell cycle state classifications. **A**. Pipeline for the quality control, training, and evaluation of the ccAFv2 classifier. Data is quality controlled and normalized using sctransform. Then the data is split 80% for training and 20% for testing using 10-fold cross validation. For each iteration three steps are performed: 1) the ANN is trained using the training dataset, 2) then the trained classifier is used to predict on the test set, and 3) the predicted labels are compared to the true labels to evaluate the accuracy of the classifier. **B**. F1 scores (integrating precision and recall, max = 1) from 10-fold cross-validation of six classification methods trained on U5-hNSCs (p-values $\leq 2.8 \times 10^{-6}$). Scores were computed for each cell cycle state across 10 testing datasets. To address class imbalance, the ccAFv2 equal class model was trained with 194 randomly selected cells per state, matching the Late G1 class, which had the fewest cells. **C**. Overlay of cell labels (O'Connor et al. 2021) and ccAFv2 predictions on the U5-hNSCs UMAP. **D**. Cell fractions of U5-hNSC defined cell labels colorized by the ccAFv2 predictions. **E**. Heatmap showing the expression levels of the top three important features for each ccAFv2 state. Data is grouped by ccAFv2-classified *in vivo* human neural stem cells collected at nine weeks post-conception (PCW 9-1) (Zeng et al, 2023)

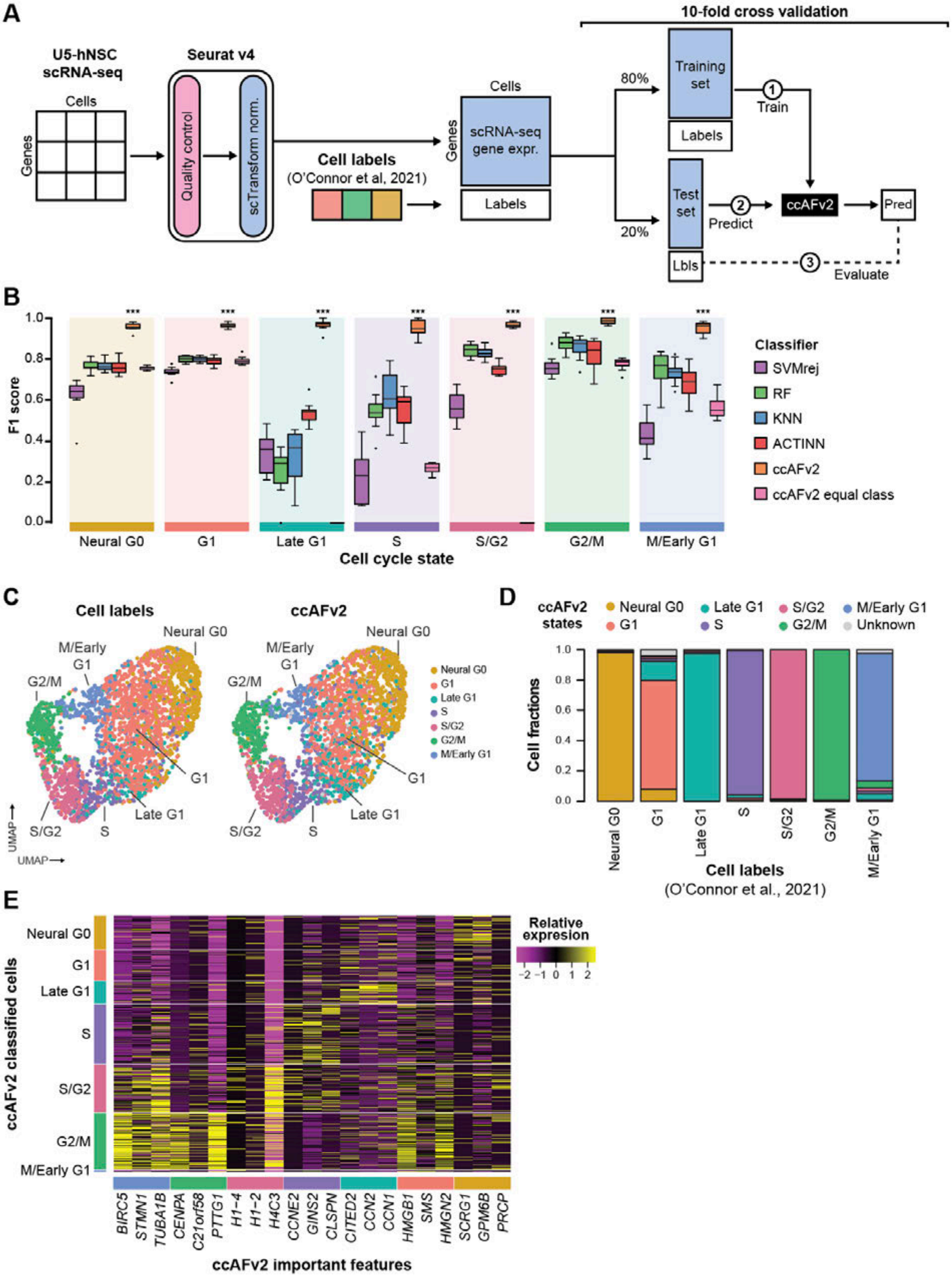
A
10-fold cross validation
U5-hNSC scRNA-seq
Cells
Genes
Seurat v4
Quality control
scTransform norm.
Cell labels
(O'Connor et al, 2021)
Cells
Genes
scRNA-seq gene expr.
Labels
80%
20%
Training set
Labels
Test set
Lbls
1 Train
2 Predict
ccAFv2
Pred
3 Evaluate
B
F1 score
Cell cycle state
Neural G0
G1
Late G1
S
S/G2
G2/M
M/Early G1
Classifier
SVMrej
RF
KNN
ACTINN
ccAFv2
ccAFv2 equal class
C
Cell labels
ccAFv2
UMAP
D
ccAFv2 states
Neural G0
G1
Late G1
S
S/G2
G2/M
M/Early G1
Unknown
Cell fractions
Cell labels
(O'Connor et al., 2021)
E
ccAFv2 classified cells
Relative expresion
BIRC5
STMN1
TUBA1B
CENPA
C21orf58
PTTG1
H1-4
H1-2
H4C3
CCNE2
GINS2
CLSPN
CITED2
CCN2
CCN1
HMGB1
SMS
HMGN2
SCRG1
GPM6B
PRCP
ccAFv2 important features

**Supplemental Figure S2.** Application of ccSeurat using the ccAFv2 marker genes to the U5-hNSCs. **A**. Venn diagram of ccSeurat S marker genes with ccAFv2 S and S/G2 marker genes. Common genes between the marker gene sets are labeled in red. **B**. Venn diagram of ccSeurat G2/M marker genes with ccAFv2 S/G2 and G2/M marker genes. Common genes between the marker gene sets are labeled in red. **C**. Overlay of cell labels (O'Connor et al. 2021) on the U5-hNSC UMAP. **D**. Overlay of ccAFv2 labels on the U5-hNSC UMAP. **E**. Overlay of the ccSeurat labels on the U5-hNSC UMAP. **F**. Overlay of the adapted (with ccAFv2 marker genes) ccSeurat labels on the U5-hNSC UMAP. The function was edited to include Neural G0, Late G1, S, S/G2, G2/M and M/Early G1 marker genes. **G**. Median AMI score for each cell cycle classifier's predictions of the U5-hNSCs relative to the U5-hNSC cell labels (O'Connor et al. 2021) is plotted against the number of cell cycle states predicted by the classifier. The average similarity to the reference was computed, based on the number of cell cycle states in the reference and predicted by the classifier, and were plotted at 10 percent intervals to facilitate comparison between classifiers with differing numbers of predicted cell cycle states.

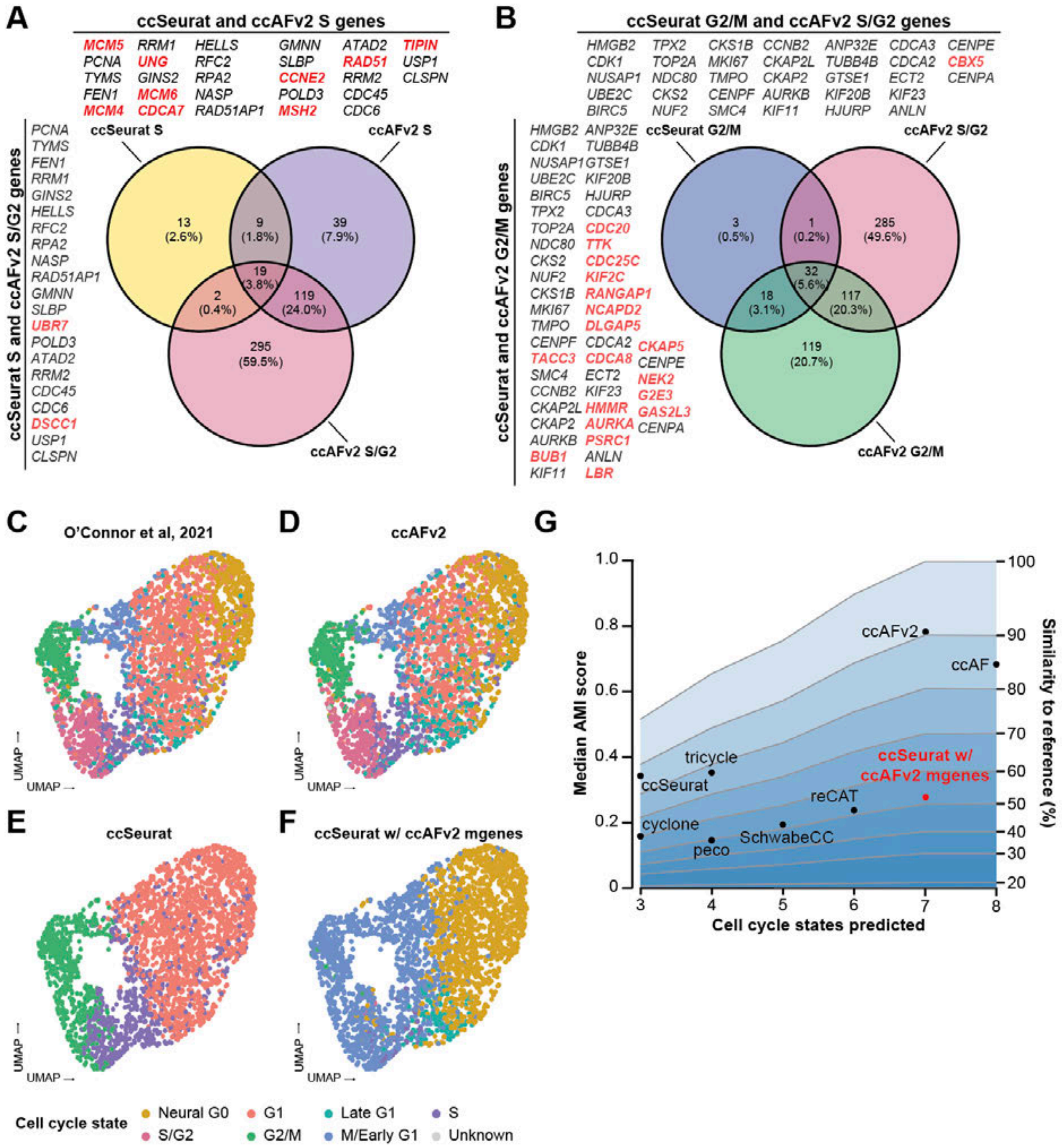
A
ccSeurat and ccAFv2 S genes
MCM5 RRM1 HELLS GMNN ATAD2 TIPIN
PCNA UNG RFC2 SLBP RAD51 USP1
TYMS GINS2 RPA2 CCNE2 RRM2 CLSPN
FEN1 MCM6 NASP POLD3 CDC45
MCM4 CDCA7 RAD51AP1 MSH2 CDC6
ccSeurat S and ccAFv2 S/G2 genes
PCNA TYMS FEN1 RRM1 GINS2 HELLS RFC2 RPA2 NASP RAD51AP1 GMNN SLBP UBR7 POLD3 ATAD2 RRM2 CDC45 CDC6 DSCC1 USP1 CLSPN
ccSeurat S
ccAFv2 S
ccAFv2 S/G2
13 (2.6%)
9 (1.8%)
39 (7.9%)
19 (3.8%)
2 (0.4%)
119 (24.0%)
295 (59.5%)
B
ccSeurat G2/M and ccAFv2 S/G2 genes
HMGB2 TPX2 CKS1B CCNB2 ANP32E CDCA3 CENPE
CDK1 TOP2A MKI67 CKAP2L TUBB4B CDCA2 CBX5
NUSAP1 NDC80 TMPO CKAP2 GTSE1 ECT2 CENPA
UBE2C CKS2 CENPF AURKB KIF20B KIF23
BIRC5 NUF2 SMC4 KIF11 HJURP ANLN
ccSeurat and ccAFv2 G2/M genes
HMGB2 ANP32E CDK1 TUBB4B NUSAP1 GTSE1 UBE2C KIF20B BIRC5 HJURP TPX2 CDCA3 TOP2A CDC20 NDC80 TTK CKS2 CDC25C NUF2 KIF2C CKS1B RANGAP1 MKI67 NCAPD2 TMPO DLGAP5 CENPF CDCA2 CKAP5 TACC3 CDCA8 CENPE SMC4 ECT2 NEK2 CCNB2 KIF23 G2E3 CKAP2L HMMR GAS2L3 CKAP2 AURKA CENPA AURKB PSRC1 BUB1 ANLN KIF11 LBR
ccSeurat G2/M
ccAFv2 S/G2
ccAFv2 G2/M
3 (0.5%)
1 (0.2%)
285 (49.6%)
32 (5.6%)
18 (3.1%)
117 (20.3%)
119 (20.7%)
C
O'Connor et al, 2021
D
ccAFv2
E
ccSeurat
F
ccSeurat w/ ccAFv2 mgenes
UMAP
UMAP
Cell cycle state
Neural G0
G1
Late G1
S
S/G2
G2/M
M/Early G1
Unknown
G
Median AMI score
Similarity to reference (%)
Cell cycle states predicted
ccAFv2
ccAF
tricycle
ccSeurat
ccSeurat w/ ccAFv2 mgenes
reCAT
cyclone
SchwabeCC
peco
1.0 0.8 0.6 0.4 0.2 0
100 90 80 70 60 50 40 30 20
3 4 5 6 7 8

**Supplemental Figure S3.** Application of ccAFv2 to hNSC in vivo and determining the optimal likelihood threshold. **A**. Cells predicted tabulated from ccAFv2 application with likelihood thresholds 0.0 to 0.9 to hNSCs from 3 – 12 PCW fetal tissue (Zeng et al. 2023). Dashed line indicates median cells predicted across all week stages. **B**. Adjusted mutual information (AMI) scores for ccAFv2 cell cycle predictions relative to the ccSeurat cell cycle states tabulated from ccAFv2 application with likelihood thresholds 0.0 to 0.9 to hNSCs from 3 – 12 PCW fetal tissue. Dashed line indicates median AMI across all week stages. Red dashed line indicates significantly improved median AMI score due to applying threshold.

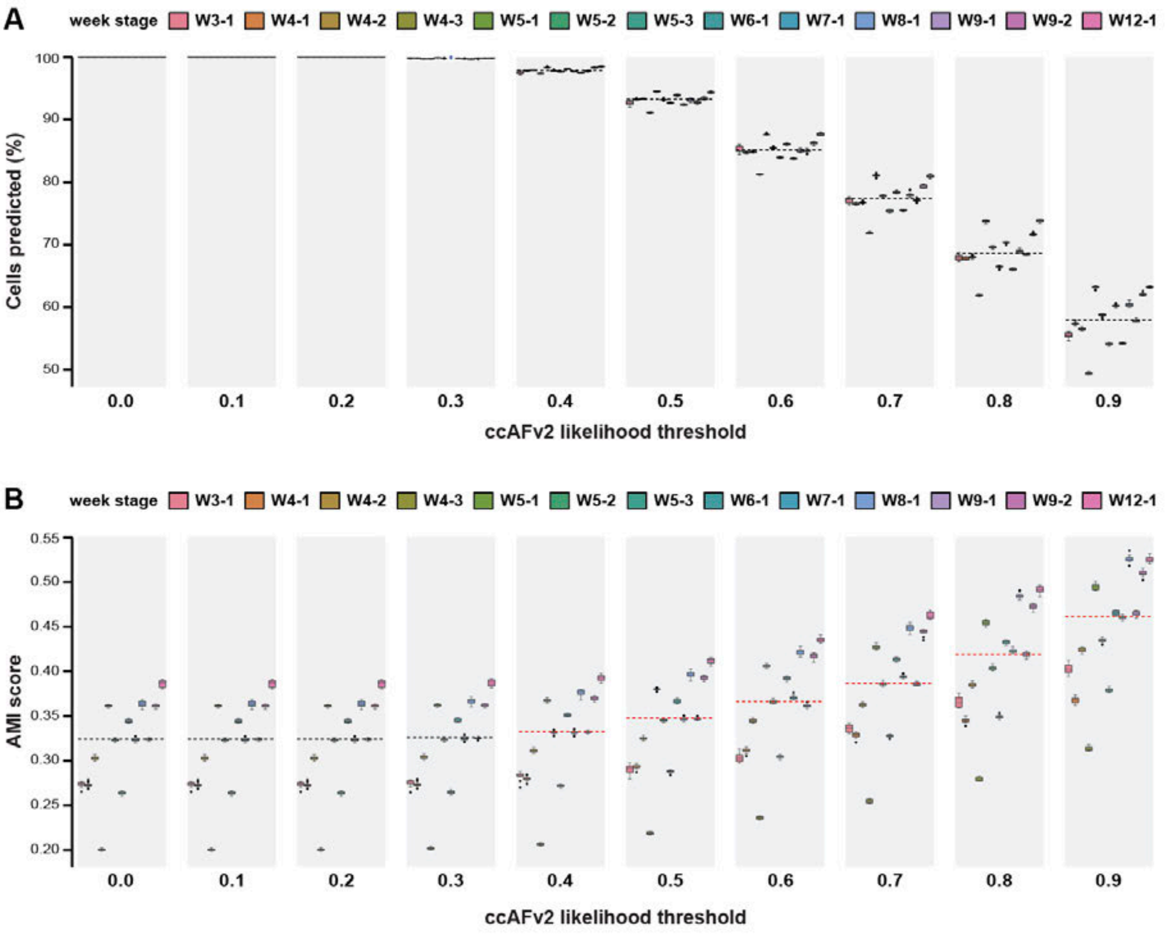

**Supplemental Figure S4.** U5-hNSC missing gene analyses. **A**. Sensitivity analysis shows how a random sampling of 10-99% information from the U5-hNSCs effects the error rate of the ccAFv2 classifier with a likelihood threshold of 0.5. **B**. Error rate in (**A**) broken down by cell cycle state. **C**. Percentage of cells predicted for U5-hNSC data broken down by cell cycle state and percentage of missing information. **D**. Sensitivity analysis shows how a random sampling of 10-99% information from the U5-hNSCs effects the error rate of the ccAFv2 classifier with a likelihood threshold of 0.7, and **E**. 0.9. **F**. ccAFv2 threshold plot for U5-hNSCs.

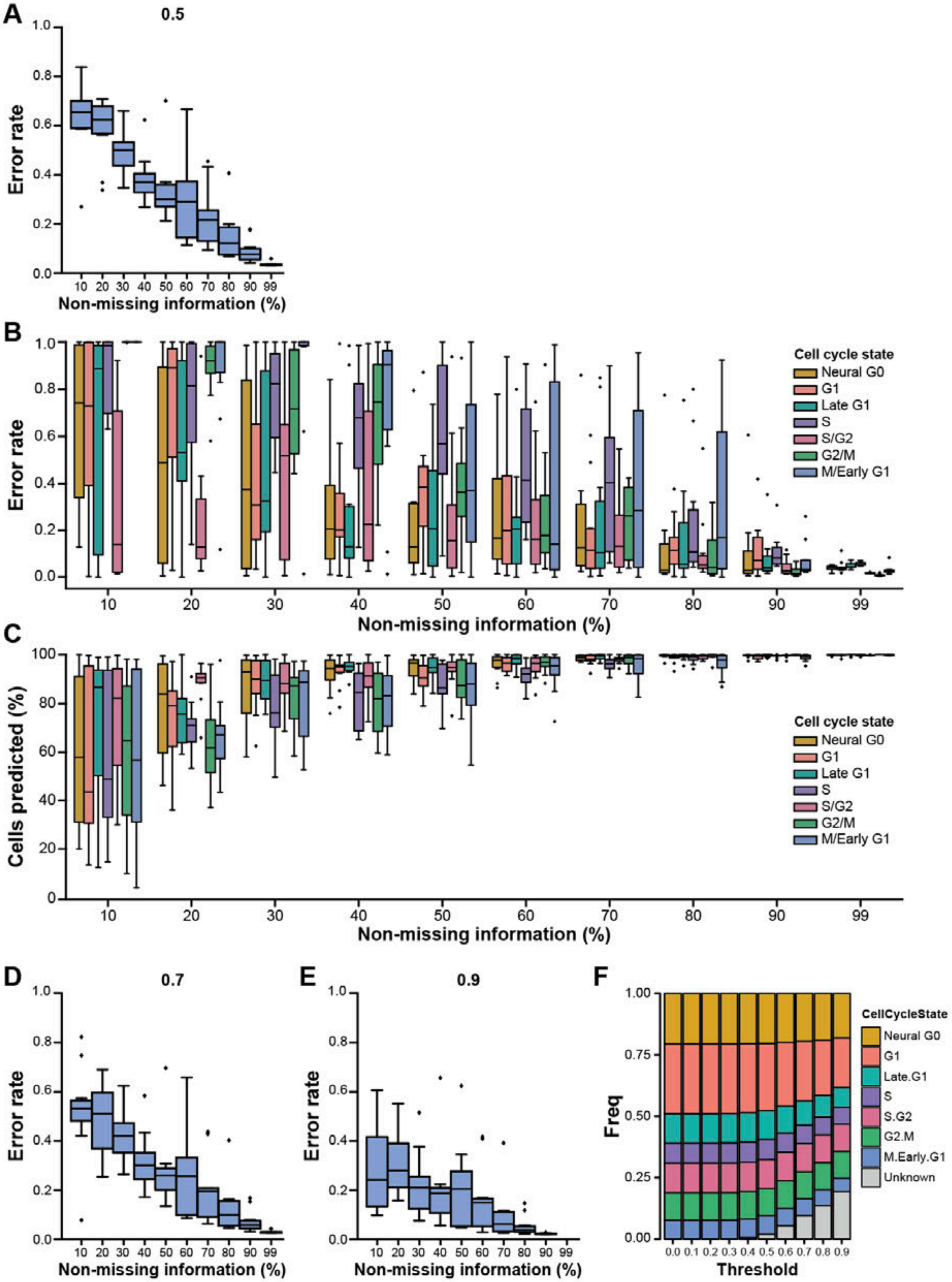

A
0.5
Error rate
Non-missing information (%)
B
Cell cycle state
Neural G0
G1
Late G1
S
S/G2
G2/M
M/Early G1
Error rate
Non-missing information (%)
C
Cells predicted (%)
Non-missing information (%)
D
0.7
E
0.9
F
CellCycleState
Neural G0
G1
Late.G1
S
S.G2
G2.M
M.Early.G1
Unknown
Freq
Threshold


302

**Supplemental Figure S5.** hNSCs from PCW8 missing gene analyses. **A.** Sensitivity analysis shows how a random sampling of 10-99% information from the PCW8-derived hNSCs from Zeng et al, 2023 effects the AMI of the ccAFv2 classifier relative to the ccSeurat predictions with a likelihood threshold of 0.5, **B.** 0.7, and **C.** 0.9.

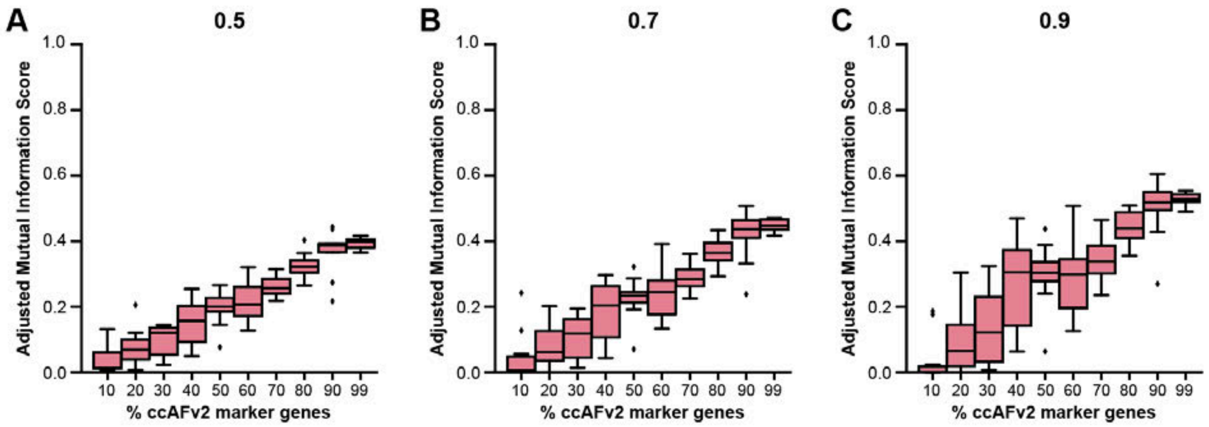

**Supplemental Figure S6.** Cyclin and CDK expression in SkMSC G0 and not G0 cells.

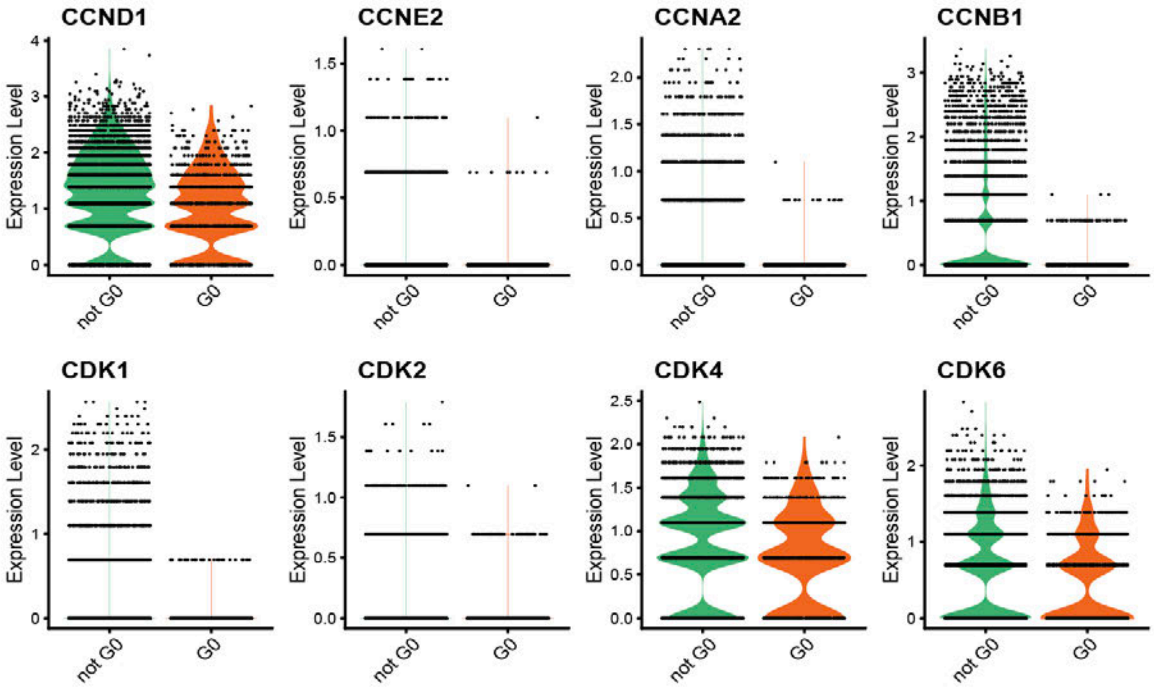

**Supplemental Figure S7**. ccAFv2 proportions for non-NSC cell types derived from fetal tissue from Zeng et al. broken up by tissue and week stage.

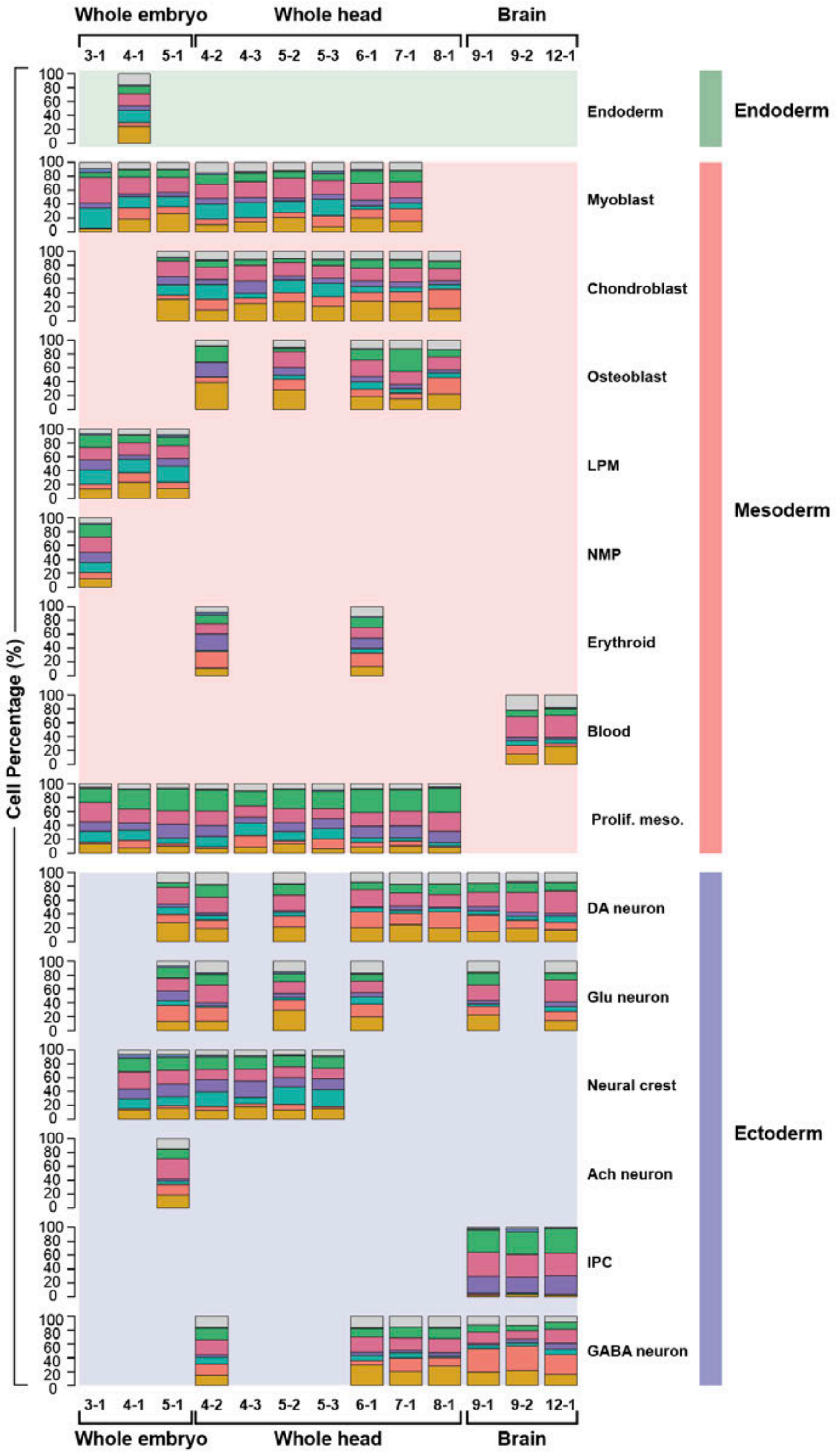

**Supplemental Figure S8.** Application of ccAFv2 to LGGs with and without growth factors. Overlay of ccAFv2 predictions on Grade 2 astrocytoma (LGG275) with growth factors (**A**) and without growth factors (**B**). **C.** Summary of the proportions of cell cycle states with and without growth factors. Overlay of ccAFv2 predictions on Grade 3 oligodendroglioma (BT237) with growth factors (**D**) and without growth factors (**E**). **F**. Summary of the proportions of cell cycle states with and without growth factors.

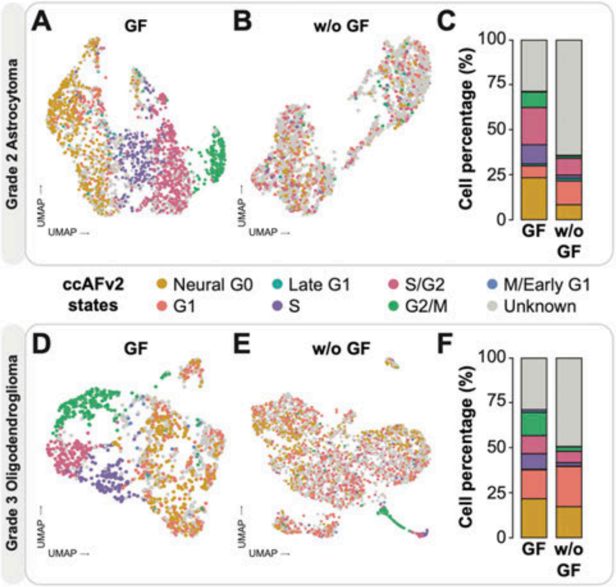

**Supplemental Figure S9.** Application of ccAFv2 to low-grade glioma scRNA-seq samples grown with and without growth factor. **A.** Cell fractions of a grade 2 astrocytoma (LGG275) ccAFv2 cell cycle classifications using likelihood thresholds 0.0 to 0.9. G = cells grown with growth factor; N = cells grown without growth factor. **B.** Cell fractions of a grade 3 oligodendroglioma (BT237) ccAFv2 cell cycle classifications using likelihood thresholds 0.0 to 0.9. G = cells grown with growth factor; N = cells grown without growth factor.

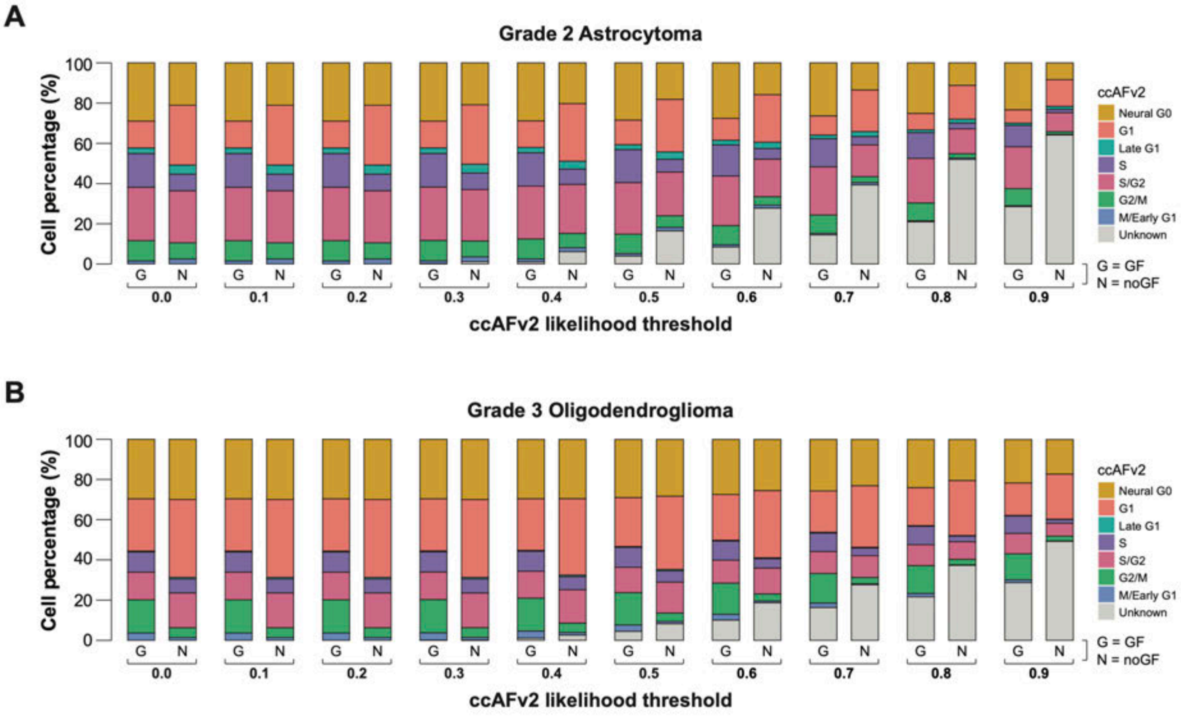

**Supplemental Figure S10.** Cell cycle regression. **A.** U5-hNSC PCA plot colorized by ccAFv2. **B.** U5-hNSC with S and G2/M ccAFv2 classes regressed PCA plot colorized by ccAFv2. **C** U5-hNSC with all cycling ccAFv2 classed regressed PCA plot colorized by ccAFv2. **D**. U5-hNSC PCA plot colorized by ccSeurat. **E**. U5-hNSC with S and G2/M ccSeurat classes regressed PCA plot colorized by ccSeurat. **F**. Venn diagram of ccAFv2 cell cycle state marker genes and ccSeurat cell cycle state marker genes.

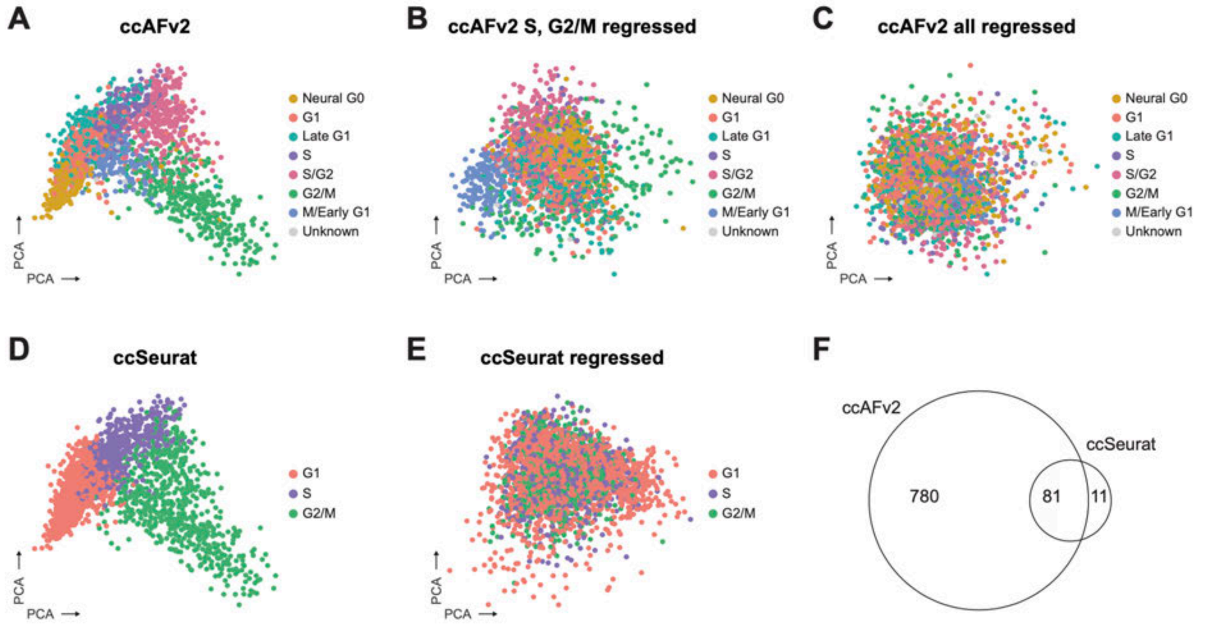

**Supplemental Figure S11.** Application of ccAFv2 to quiescent (qNSC) and activated (aNSC) neural stem cells from Llorens-Bobadilla et al, 2015. **A.** Cell fractions of qNSC1 ccAFv2 cell cycle classifications using likelihood thresholds 0.0 to 0.9. **B.** Cell fractions of qNSC2 ccAFv2 cell cycle classifications using likelihood thresholds 0.0 to 0.9. **C.** Cell fractions of aNSC1 ccAFv2 cell cycle classifications using likelihood thresholds 0.0 to 0.9. **D.** Cell fractions of aNSC2 ccAFv2 cell cycle classifications using likelihood thresholds 0.0 to 0.9.

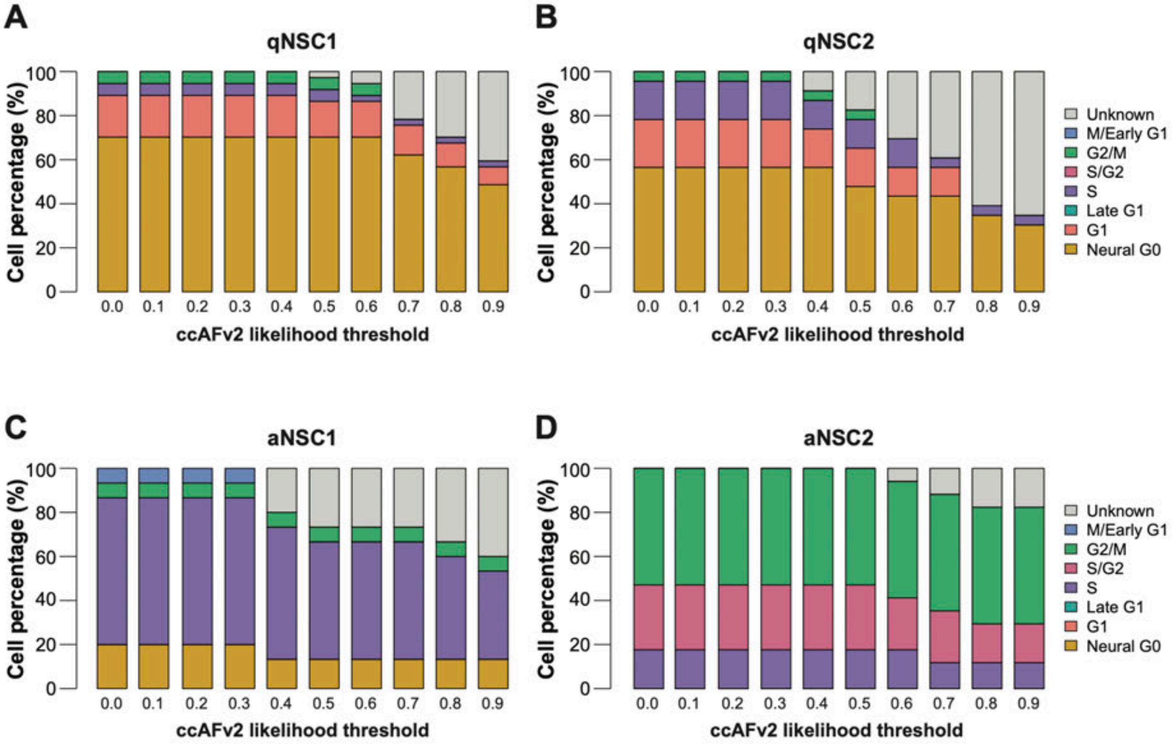

**Supplemental Figure S12.** Validation of cell cycle states in U5-hNSC data using QuieScore and cyclin expression levels. **A**. Application of QuieScore to the U5-hNSCs (q_score_raw > 3 defined G0 cells). **B**. Hypergeometric enrichment of QuieScore and U5-hNSC cell labels (O’Connor et al, 2021). Values reported are the negative log10 of the p-values.

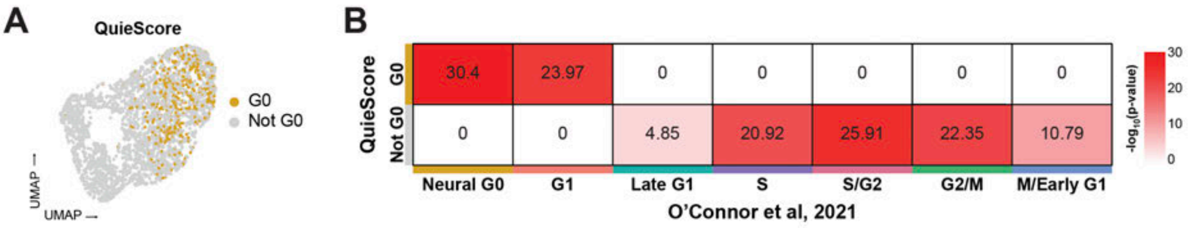

**Supplemental Figure S13.** Expression of skin and bone marker genes to the developing cortex of a male C57BL/6 mouse embryo at E15.5. **A**. H&E staining for the developing embryo cortex. **B**-**C**. Expression of skin (*Krt5*) and bone (*Col1a1*) key marker genes in the developing embryo cortex.

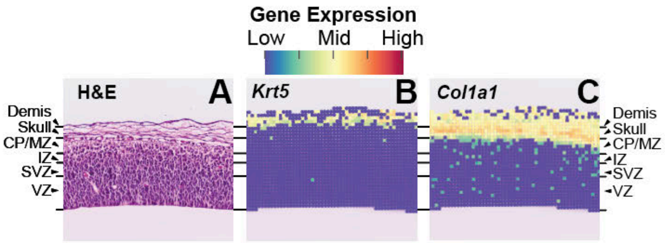

**Supplemental Figure S14.** Application of the R and python-based ccAFv2. **A.** Cell percentages of *in vivo* hNSCs from PCW8 from Zeng et al, 2023 of ccAFv2 cell cycle state classifications using the R package Seurat v4 and v5, and Python package Scanpy.

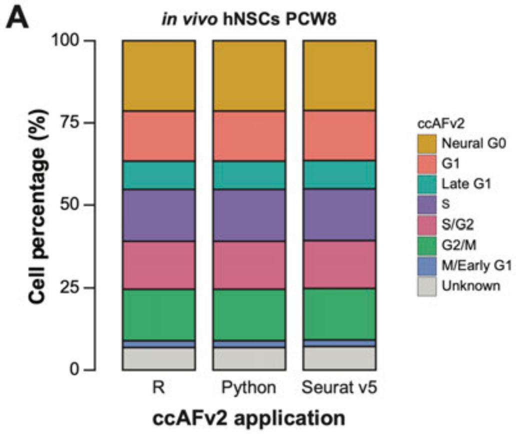
